# The schizophrenia-associated variant in *SLC39A8* alters N-glycosylation in the mouse brain

**DOI:** 10.1101/2020.12.22.424076

**Authors:** Robert G. Mealer, Sarah E. Williams, Maxence Noel, Bo Yang, Alexandria D’Souza, Toru Nakata, Daniel B. Graham, Elizabeth A. Creasey, Murat Cetinbas, Ruslan Sadreyev, Edward M. Scolnick, Christina M. Woo, Jordan W. Smoller, Ramnik J. Xavier, Richard D. Cummings

## Abstract

A missense mutation (A391T) in the manganese transporter *SLC39A8* is strongly associated with schizophrenia in genomic studies, though the molecular connection to the brain remains hypothetical. Human carriers of A391T have reduced serum manganese, altered plasma glycosylation, and brain MRI changes consistent with altered metal transport. Here, using a knock-in mouse model homozygous for A391T, we show that the schizophrenia-associated variant changes protein glycosylation in the brain. N-linked glycosylation was most significantly impaired, with effects differing between regions. RNAseq analysis showed negligible regional variation, consistent with changes in the activity of glycosylation enzymes rather than gene expression. Finally, nearly one third of detected glycoproteins were differentially N-glycosylated in the cortex, including members of several pathways previously implicated in schizophrenia such as cell adhesion molecules and neurotransmitter receptors. These findings provide a mechanistic link between a risk allele and biochemical changes in the brain, furthering our molecular understanding of the pathophysiology of schizophrenia.

## Introduction

Psychiatric disorders are heterogeneous in nearly every aspect - symptoms, diagnosis, treatment, and course - but a common feature is the lack of clear disease mechanisms to guide therapeutic development. Schizophrenia is a diagnostic classification for individuals afflicted with a combination of cognitive dysfunction, social withdrawal, functional regression, and psychotic symptoms^1^. It affects between 0.5 - 1.0% of the population and can lead to a lifetime of significant disability, economic and social hardships, increased mortality and decreased life expectancy^2^. Age of onset is often during adolescence and early adulthood, suggesting changes in brain development and maturation play a critical role^3^. Studies of disease mechanisms in schizophrenia were jumpstarted by the observation that dopamine antagonists provide some level of symptomatic relief, which led to decades of research on neurotransmitter signaling^4–6^. Despite this progress, no treatment with a novel mechanism of action has been approved in nearly 50 years.

Genetic liability is the strongest epidemiologic contributor to overall risk for schizophrenia^7^. Current evidence for schizophrenia and most psychiatric disorders is consistent with a polygenic inheritance pattern, with disease contributions from many common variants of small effect and few rare mutations of large effect^8–10^, originally hypothesized over fifty years ago^11^. A major advance came in 2014 with the publication of over 100 loci associated with schizophrenia through GWAS^12^, with the most recent studies implicating 270 common loci^13^. Most associated variants are in the non-coding region of the genome, requiring careful follow-up studies to map the affected gene and pathophysiologic consequence. The first successful example of such a study involves the complement component 4 (C4) genes (*C4A* and *C4B*) near the MHC locus^14^. Individuals with schizophrenia have increased expression of brain *C4A*, and the schizophrenia-associated allele increases *C4A* expression in the brain. Global deletion of the *C4* gene in mice, which corresponds to both *C4A* and *C4B* genes in humans, leads to decreased complement deposition and synaptic pruning during post-natal development^14^, providing a potential link between a locus associated with schizophrenia and a molecular pathway implicated in schizophrenia pathogenesis^15^.

For the remaining loci, understanding their connection to schizophrenia is a considerable challenge, particularly for those without clear roles in the brain. One such example is the group of genes involved in protein glycosylation, the post- and co-translational attachment of carbohydrate structures to asparagine (N-linked) or serine, threonine and tyrosine (O-linked) residues, respectively^16,17^. In addition to at least 5 glycosylation enzymes, a variant in the manganese (Mn^2+^) transporter *SLC39A8* is strongly associated with schizophrenia^12,18^. A large number of glycosyltransferases require Mn^2+^ as an obligatory co-factor^19–21^, and numerous case series have described congenital disorders of glycosylation (CDG) caused by a near total absence of Mn^2+^ in severe homozygous mutation carriers in *SLC39A8*^22–25^. *SLC39A8* is expressed at relatively low levels in most tissues of the body including the brain^26,27^, and single-cell sequencing data from mouse brain suggest the transporter is enriched within endothelial cells^28^. The schizophrenia-associated variant in *SLC39A8* of the single-nucleotide polymorphism rs13107325 is of particular interest, as it corresponds to a missense mutation (A391T) recently confirmed as the causal variant within *SLC39A8*^13^. A391T is present in ~8% of the general population, with increased prevalence in European lineages and near absence in Asian and African lineages^29^. In addition to its role in schizophrenia, rs13107325 is one of the most pleotropic variants in the genome and is associated with dozens of traits through GWAS^30^, including several neuropsychiatric phenotypes and a decrease in serum Mn^2+^ levels^31–35^. Like other common variants, the relative effect of rs13107325 on most phenotypes is small, e.g., reducing the concentration of serum Mn^2+^ by ~10%^18,35^ and increasing the odds ratio of schizophrenia from 1.0 to 1.15 in heterozygous carriers^12,13^.

The pathophysiologic connection between A391T and most conditions remain hypothetical^29,36–39^. We recently described serum glycosylation changes in homozygous *SLC39A8* mutation carriers who presented with symptoms of a CDG but who were missed by conventional diagnostic methods. Subtle dysglycosylation differences were observed in heterozygous carriers as well, consistent with a dominant effect of SLC39A8-CDG alleles^40^. Lin and colleagues describe an increase of one precursor N-glycan in plasma from a small number of homozygous A391T carriers, suggestive of hypogalactosylation^41^. We subsequently reported a detailed N-glycomic analysis of plasma glycoproteins from homozygous and heterozygotes A391T carriers, and identified dysglycosylation in both genotypes characterized by reduced complexity and branching of large N-glycans, primarily those synthesized by the liver^42^. In that study, we also described a previously unappreciated sex-dependent effect of the A391T variant on glycosylation, with males showing a larger alteration in plasma N-glycosylation compared to females^42^. Although plasma glycosylation is an important peripheral marker for enzymatic activity and disease state in some scenarios, it is unlikely to reflect changes in the brain. However, analysis of brain MRI data from the UK Biobank of A391T mutation carriers identified a difference in the T2w:T1w ratio in several regions, consistent with an effect of A391T on brain metal transport^42^.

Here, to further understand the molecular role of the schizophrenia-associated variant in the brain, we utilized a murine model expressing the homologous human mutation A391T in murine *Slc39a8* (A393T)^43^. By applying several methods optimized for analysis of protein glycosylation in the brain^44^, in addition to glycoproteomics and a novel method for glycan quantification, we present a comprehensive portrait of dysglycosylation in the adult A391T homozygous mouse brain. These results are evidence of a biochemical change in the brain directly caused by a schizophrenia risk variant.

## Results

### A391T changes brain N-glycosylation in a region-specific manner

We utilized a recently described mouse model expressing the schizophrenia-associated human variant A391T in *SLC39A8* within the murine gene (Ala 393 in *Slc39a8*), which exhibits no overt neurobehavioral or developmental phenotype and reproduces at the expected frequencies^43^. We hypothesized that A391T would decrease Mn^2+^ concentration and subsequent activity of Mn^2+^-dependent Golgi glycosyltransferases in the brain. The N-glycome of the brain is unique compared to other organs - consisting predominantly of high-mannose, bisected and fucosylated glycans, with a much lower abundance of complex, branched structures containing galactose and sialic acid - with subtle but significant differences between brain regions and sexes^44^.

We first analyzed the effect of A391T on the relative abundance of N-glycans in four independent brain regions (frontal cortex, hippocampus, striatum and cerebellum) of male and female adult mice with semi-quantitative MALDI-TOF N-glycomics following Peptide N-Glycosidase (PNGase F) release and permethylation. Homozygous mutant mice (TT) were compared to wild-type littermate controls (CC) at 12-weeks of age (**Table 1**). We identified bidirectional and region-specific changes of the A391T mutation on N-glycosylation, with a larger effect consistently observed in males (**Fig. 1A**, **Fig. S1**). N-glycans appeared more complex in the male TT cortex compared to CC controls, with greater relative abundance of tri- and tetra-antennary structures along with a corresponding decrease in high-mannose glycans (**Fig. 1B**). Glycans containing fucose, galactose, and sialic acid were also increased in TT cortex, consistent with an apparent increase in complexity of N-glycans. In the hippocampus, which is evolutionarily most related to the cortex^45^, the pattern of N-glycosylation changes in TT mice were similar though did not reach statistical significance, while N-glycans in the striatum appeared mostly unaffected. In contrast, the cerebellum of TT mice displayed less complex N-glycans compared to CC controls, with a significant reduction in sialylated structures (**Fig 1B**). Several individual N-glycans differed significantly between genotypes, the majority of which were in the cortex and cerebellum (**Table S1**). Full descriptions of individual N-glycan characteristics and abundance by region and sex are provided in supplementary material (**Table S1, S2**). Analysis of N-glycans from cortex and cerebellum at 4-weeks of age in a separate cohort failed to detect any difference between CC and TT mice (**Table S3**).

**Table 1.**
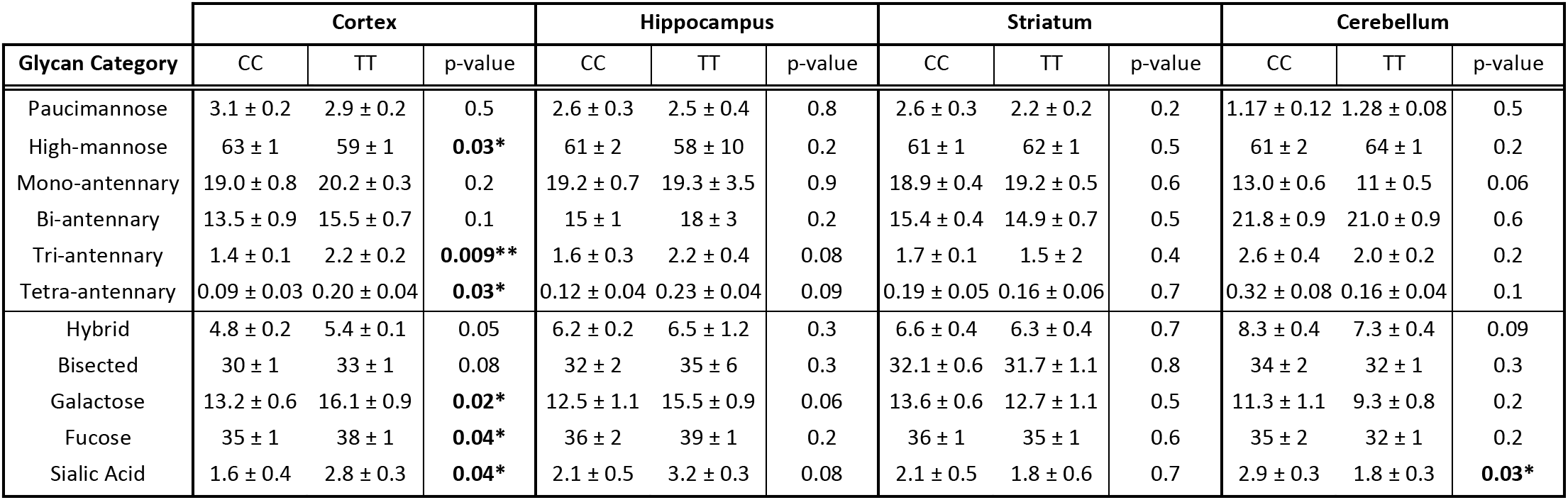
Categorical analysis of protein N-glycans revealed region-specific changes in male A391T brain. Data presented as mean percent abundance +/− SEM. Unpaired t-tests performed for each category assuming unequal variance.

| Glycan Category | Cortex |  |  | Hippocampus |  |  | Striatum |  |  | Cerebellum |  |  |
| --- | --- | --- | --- | --- | --- | --- | --- | --- | --- | --- | --- | --- |
|  | CC | TT | p-value | CC | TT | p-value | CC | TT | p-value | CC | TT | p-value |
| Paucimannose | 3.1 ± 0.2 | 2.9 ± 0.2 | 0.5 | 2.6 ± 0.3 | 2.5 ± 0.4 | 0.8 | 2.6 ± 0.3 | 2.2 ± 0.2 | 0.2 | 1.17 ± 0.12 | 1.28 ± 0.08 | 0.5 |
| High-mannose | 63 ± 1 | 59 ± 1 | **0.03\*** | 61 ± 2 | 58 ± 10 | 0.2 | 61 ± 1 | 62 ± 1 | 0.5 | 61 ± 2 | 64 ± 1 | 0.2 |
| Mono-antennary | 19.0 ± 0.8 | 20.2 ± 0.3 | 0.2 | 19.2 ± 0.7 | 19.3 ± 3.5 | 0.9 | 18.9 ± 0.4 | 19.2 ± 0.5 | 0.6 | 13.0 ± 0.6 | 11 ± 0.5 | 0.06 |
| Bi-antennary | 13.5 ± 0.9 | 15.5 ± 0.7 | 0.1 | 15 ± 1 | 18 ± 3 | 0.2 | 15.4 ± 0.4 | 14.9 ± 0.7 | 0.5 | 21.8 ± 0.9 | 21.0 ± 0.9 | 0.6 |
| Tri-antennary | 1.4 ± 0.1 | 2.2 ± 0.2 | **0.009\*\*** | 1.6 ± 0.3 | 2.2 ± 0.4 | 0.08 | 1.7 ± 0.1 | 1.5 ± 2 | 0.4 | 2.6 ± 0.4 | 2.0 ± 0.2 | 0.2 |
| Tetra-antennary | 0.09 ± 0.03 | 0.20 ± 0.04 | **0.03\*** | 0.12 ± 0.04 | 0.23 ± 0.04 | 0.09 | 0.19 ± 0.05 | 0.16 ± 0.06 | 0.7 | 0.32 ± 0.08 | 0.16 ± 0.04 | 0.1 |
| Hybrid | 4.8 ± 0.2 | 5.4 ± 0.1 | 0.05 | 6.2 ± 0.2 | 6.5 ± 1.2 | 0.3 | 6.6 ± 0.4 | 6.3 ± 0.4 | 0.7 | 8.3 ± 0.4 | 7.3 ± 0.4 | 0.09 |
| Bisected | 30 ± 1 | 33 ± 1 | 0.08 | 32 ± 2 | 35 ± 6 | 0.3 | 32.1 ± 0.6 | 31.7 ± 1.1 | 0.8 | 34 ± 2 | 32 ± 1 | 0.3 |
| Galactose | 13.2 ± 0.6 | 16.1 ± 0.9 | **0.02\*** | 12.5 ± 1.1 | 15.5 ± 0.9 | 0.06 | 13.6 ± 0.6 | 12.7 ± 1.1 | 0.5 | 11.3 ± 1.1 | 9.3 ± 0.8 | 0.2 |
| Fucose | 35 ± 1 | 38 ± 1 | **0.04\*** | 36 ± 2 | 39 ± 1 | 0.2 | 36 ± 1 | 35 ± 1 | 0.6 | 35 ± 2 | 32 ± 1 | 0.2 |
| Sialic Acid | 1.6 ± 0.4 | 2.8 ± 0.3 | **0.04\*** | 2.1 ± 0.5 | 3.2 ± 0.3 | 0.08 | 2.1 ± 0.5 | 1.8 ± 0.6 | 0.7 | 2.9 ± 0.3 | 1.8 ± 0.3 | **0.03\*** |

**Fig. 1.**
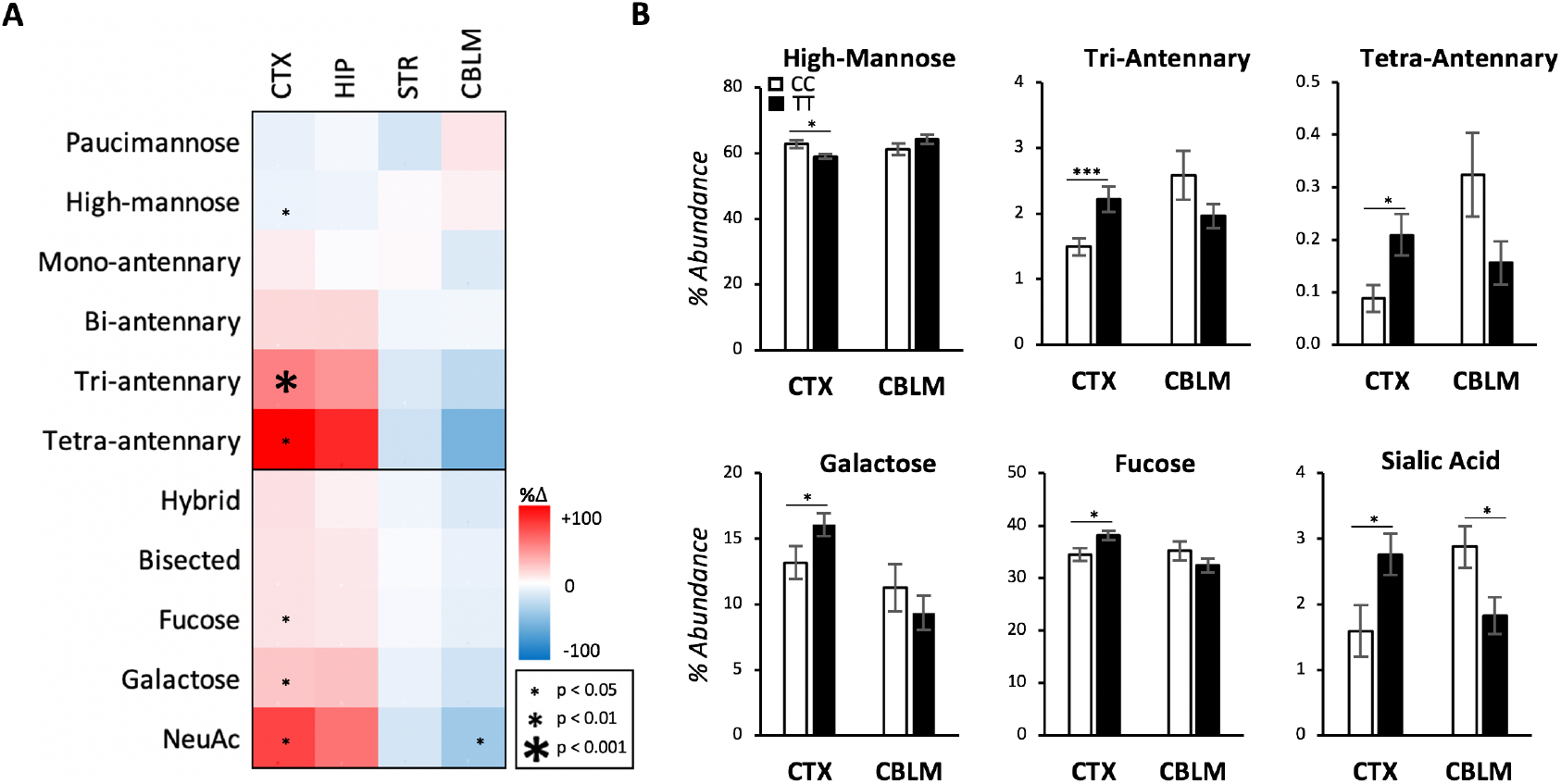
A391T alters the relative abundance of N-glycans in a region-dependent manner. A) Categorical analysis of N-glycans from the cortex (CTX), hippocampus (HIP), striatum (STR), and cerebellum (CBLM) of male mice revealed a bi-directional effect of the A391T variant. Data presented as a heat map of percent change in glycan abundance comparing TT mice to CC controls. T = 6, CC = 8 for each region. B) TT cortex showed an increase in complex N-glycans, while the cerebellum showed a downward trend in the same categories. Data presented as mean percent abundance +/− SEM.

### O-Mannose glycans are increased in A391T cortex

We are unaware of previous studies of O-glycosylation in relation to SLC39A8. After PNGase F treatment, O-glycans were removed from glycoproteins through β-elimination, which represents glycans released from serine and threonine residues, followed by permethylation and MALDI-TOF MS. This generated interpretable O-glycan spectra with 26 unique O-glycans in a smaller subset compared to our analysis of N-glycosylation (**Table S4)**. Across the four brain regions, only TT cortex showed a significant increase in the relative abundance of O-mannose (O-Man) to O-GalNAc structures (**Fig S2, Table S5**). The majority of O-glycosylation features were unchanged, with only small differences noted in structures of low abundance (cerebellar NeuGc content), or the group with the smallest sample size (hippocampus). A greater proportion of O-Man was also present in the small number of female TT cortex samples analyzed, though this difference fell short of significance (**Fig S3**).

### Total N-glycan concentration is decreased in A391T cortex

MALDI-TOF MS provides a semi-quantitative analysis following normalization within a sample. For example, if the absolute concentration of every glycan was uniformly reduced by 50%, but the relative abundance of each glycan remained the same, such a change would be undetectable using standard MALDI-TOF MS techniques. As A391T appeared to have opposite effects on the relative abundance of complex glycans in the cortex and cerebellum, we explored approaches for absolute N-glycan measurements.

Glycoprotein blotting using the lectin Concanavalin A (ConA), which has an affinity for the core α-mannose structure present in most N-glycans, was decreased ~10% in both cortex and cerebellum but fell short of significance (**Fig S4**). This was not surprising given the limited sensitivity of quantitative western blotting, particularly for a change predicted to be small from a common genetic variant. As such, we sought to develop a more sensitive measure for N-glycan concentrations within biological samples. To this end, following release with PNGase F, N-glycans were derivatized at their reducing ends with the fluorescent-linker F-MAPA^46^, resulting in addition of one fluorescent molecule to each N-glycan (**Fig 2A**). After purification, derivatized N-glycans from the isolated glycoprotein fetuin, as well as mouse serum and brain, produced linear fluorescent signals across a broad range of protein concentrations, highlighting the quantitative utility of this assay across several tissues (**Fig S4**).

**Fig. 2.**
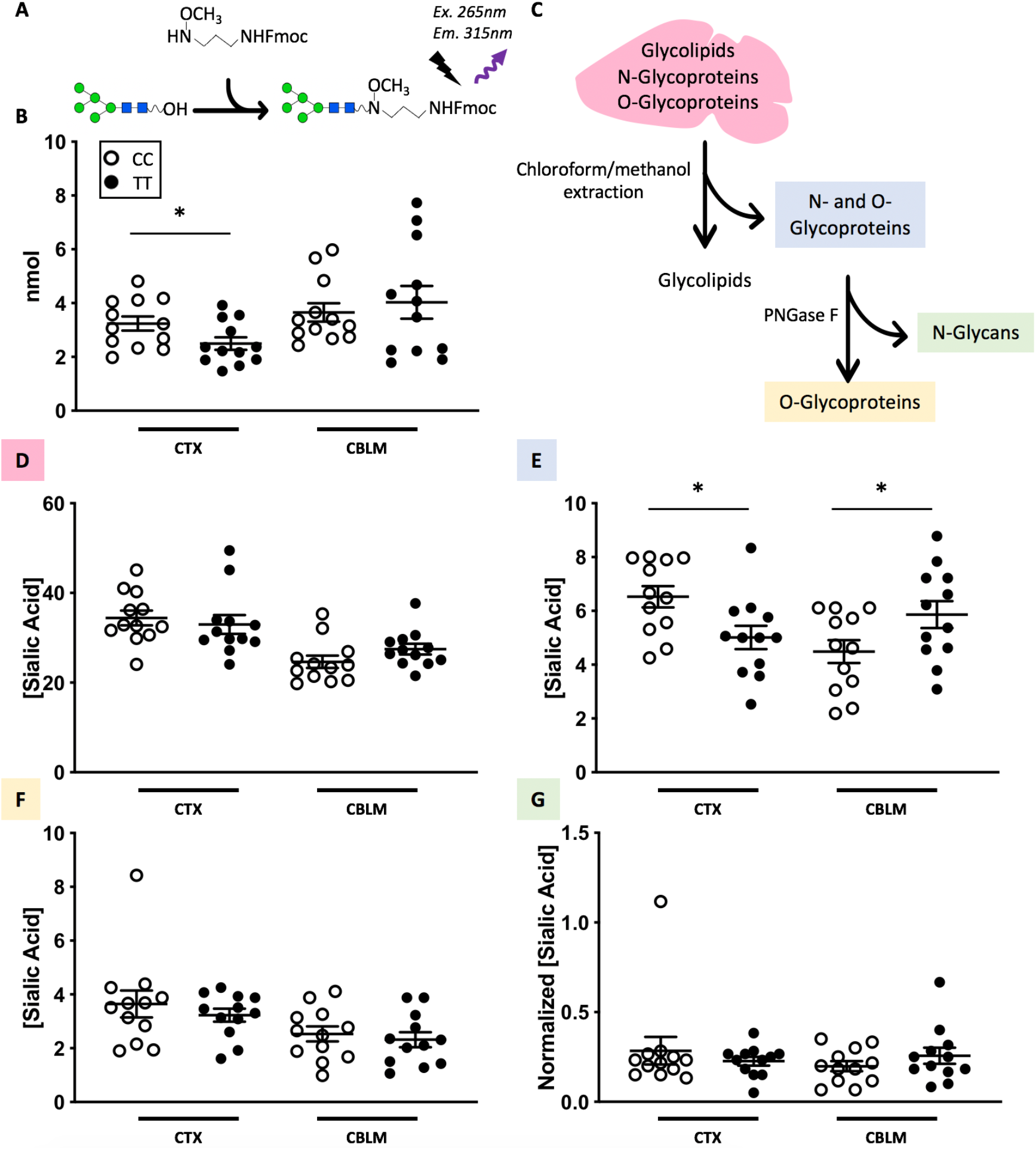
A391T lowers the absolute quantity of N-glycans in the cortex. A) Schematic of F-MAPA fluorescent glycan derivatization. Free hydroxyl groups of PNGase F-cleaved N-glycans are labeled in a 1:1 ratio with F-MAPA and quantified using fluorescence (Excitation: 265nm; Emission: 315nm). B) Quantification of brain N-glycans with F-MAPA derivatization, showing a significant decrease in the cortex of male A391T mice. C) Workflow for isolation of different glycan fractions from brain homogenate for sialic acid quantification. D) Sialic acid quantification of total lysate (glycolipids and glycoproteins), showing no difference in A391T mice. E) Sialic acid quantification of total glycoproteins (N- and O-), showing a decrease in cortex and increase in cerebellum of A391T mice. F) Sialic acid quantification of O-glycoproteins, showing no difference in A391T mice. G) Sialic acid quantification of released N-glycans, showing now difference in A391T mice when normalizing for per nmol of N-glycans. Sialic acid concentration reported as nmol sialic acid/mg of protein. N = 4 male mice per group, measured in triplicate. Individual data points are shown, with brackets representing group means +/− SEM.

Measurement of derivatized N-glycans with F-MAPA showed a decrease in the cortex of TT mice, indicating reduced absolute concentration of N-glycans (**Fig 2B**). To address the discrepancy between the semi-quantitative increase in highly branched/sialylated N-glycans identified by glycomics and the fully quantitative decrease in total N-glycans in TT cortex, we quantified total sialic acid in glycoconjugates in different brain fractions between genotypes (**Fig 2C**). Consistent with prior studies^47^, the bulk of sialic acid in both cortex and cerebellum was contained in glycolipids and removed by extraction with chloroform and methanol (**Fig S5**). The remaining sialic acid is divided roughly in half between glycoprotein N- and O-glycans based on PNGase F release of total N-glycans. These results, in combination with our glycomics observation that only a small fraction of the N-glycan pool is sialylated (~2%, compared to 90% of O-glycans)^44^, indicate that N-glycans are several-fold more abundant in the brain relative to O-glycans. No difference in sialic acid concentration was detected in total brain lysate (glycolipids and glycoproteins) between CC and TT genotypes (**Fig 2D**). After removal of glycolipids, the amount of sialic acid in the glycoprotein fraction was reduced in TT cortex and increased in TT cerebellum (**Fig 2E**). This difference was eliminated with PNGase F treatment (**Fig 2F**), indicating that the difference in amounts of sialic acid concentration was specific to N-glycans. The differing sialic acid amounts could be explained by either a change in the number of sialic acid monosaccharides per glycan, or a change in the absolute number of glycans present. When normalized for glycan concentration using results from F-MAPA-derivatized N-glycans of the same sample, we detected no difference between region or genotype in the amount of sialic acid per N-glycan (**Fig 2G**). Taken together, these results demonstrate that the absolute pool of N-glycans was decreased in the cortex of TT mice, while the relative amount of sialylated N-glycans, a minor fraction of all N-glycans, was increased in response to the A391T mutation.

### Differences in gene expression do not account for glycosylation changes in A391T cortex

We employed global RNAseq to determine if changes in gene expression contributed to glycosylation differences observed in A391T cortex and cerebellum. Minimal differences in gene expression were detected among the ~14,000 transcripts measured between CC and TT cortex (**Fig 3A**) and cerebellum (**Fig 3B**) by RNAseq, with only 3 and 42 transcripts being differentially expressed on a transcriptome-wide level, respectively (**Table S6**). Of note, there was no significant difference in expression of glycosyltransferases, glycosylhydrolases, or *Slc39a8* in either region between genotypes. Comparison between cortex and cerebellum within each genotype (**Fig 3C**, **Fig 3D**) showed nearly identical regional expression patterns, consistent with known regional differences (**Fig 3E**, **Fig S6**).

**Fig. 3.**
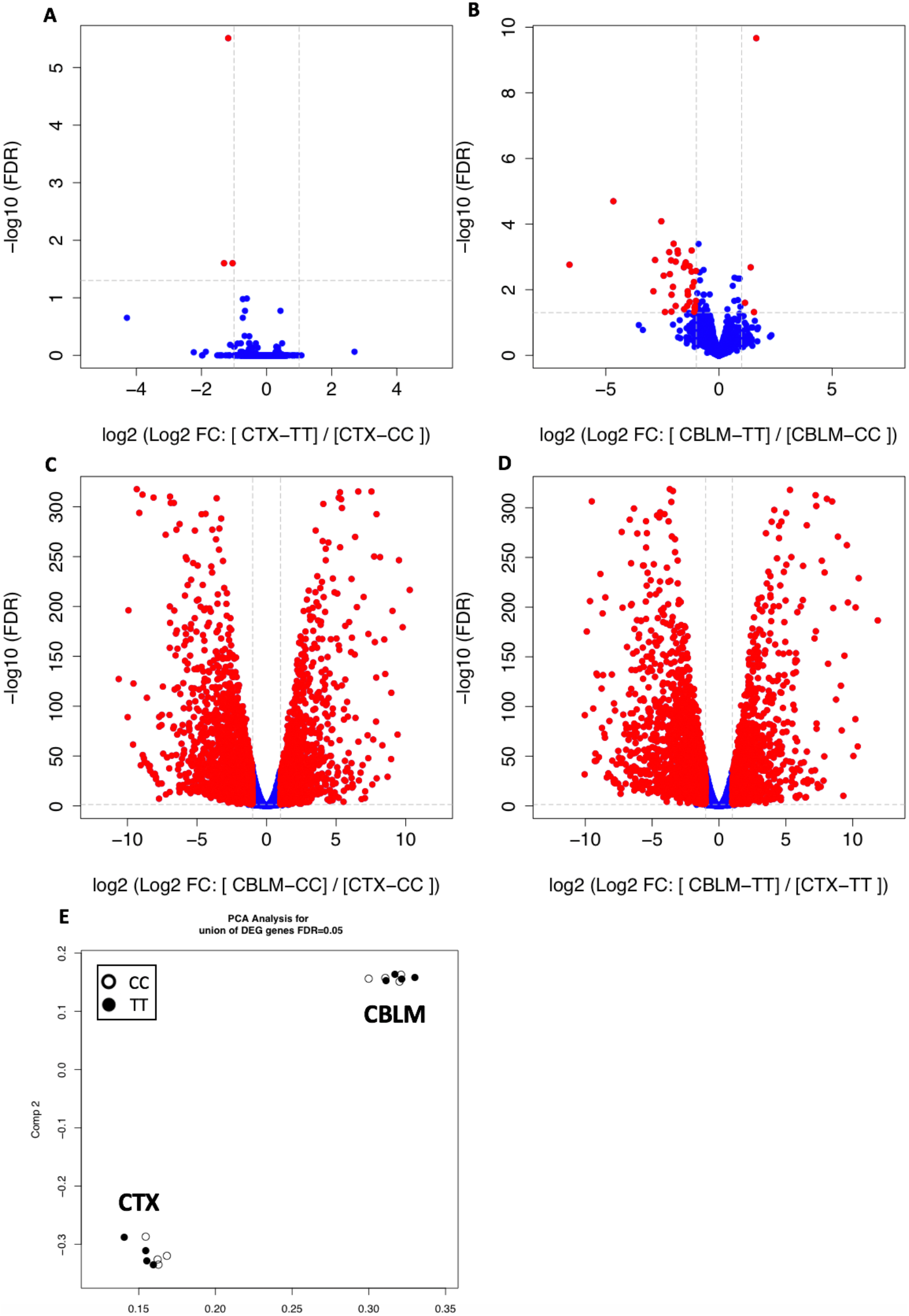
RNAseq analysis identifies minimal gene expression changes in the cortex and cerebellum of A391T mice. Volcano plots comparing cortex (A) and cerebellum (B), identifies minimal gene expression changes in A391T mice based on genotype. Volcano plots between cortex and cerebellum in both wild-type (C) and A391T (D) mice are consistent with known regional differences in gene expression. E) PCA analysis highlights the lack of gene expression differences between genotypes in A391T mice, with CC (white dots) and TT (black dots) from cortex and cerebellum shown. N = 4 male mice per group.

### N-glycoprotein levels are altered in A391T cortex

Glycomic and RNAseq data were consistent with changes in the enzymatic activity of glycosyltransferases in A391T mice rather than their expression. However, this does not provide information on altered target protein glycosylation, which could be affected in a broad or target-specific manner. To obtain quantitative data on a protein-specific level, we used hydrophilic interaction liquid chromatography (HILIC) and multi-lectin affinity enrichment to isolate tryptic N-glycopeptides from the cortex of male CC and TT mice, followed by analysis with LC-MS/MS. A total of 517 peptide sequence fragments (PSMs) with 554 unique N-glycosylation consensus sites were detected, originating from 210 distinct glycoproteins (**Table S7**). Near uniform N-glycoprotein levels were seen in the five CC samples, and TT mice showed a similar pattern in 4 of 5 samples, with one sample appearing as an outlier from both CC and TT groupings (**Fig 4A, B**). Repeat genotyping confirmed that both homozygous mutation status and sex were accurate, and the sample was included in subsequent analyses. Of the 210 N-glycoproteins, 66 had significantly different levels in TT cortex, with 41 decreased and 25 increased based on an adjusted p value <0.05 and fold change > 2.0 (**Fig 4C**). The full N-glycoproteomic data can be found in the supplemental data and via ProteomeXchange with identifier PXD021632.

**Fig. 4.**
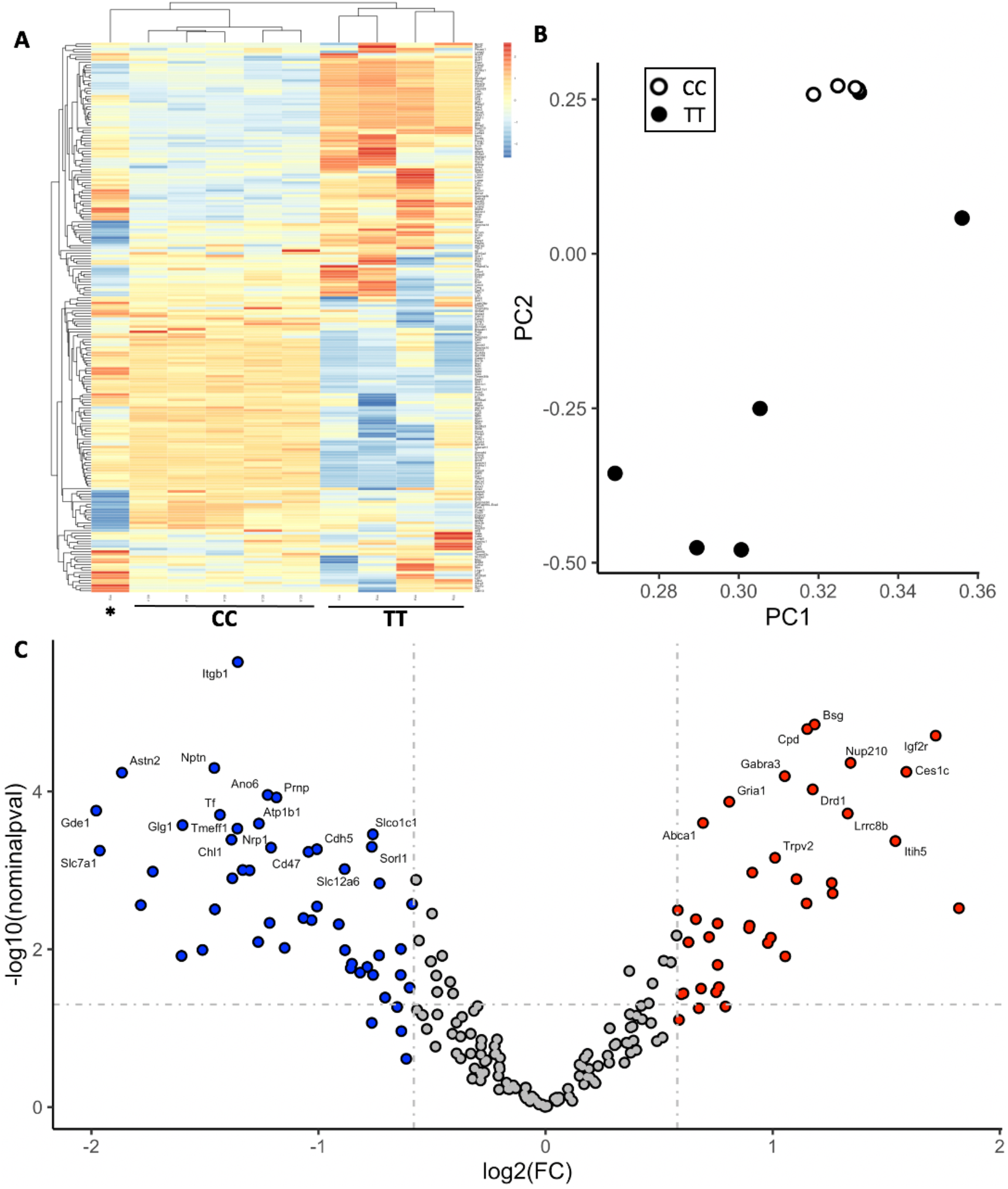
A391T mouse cortex has altered N-glycosylation of one third of glycoproteins. A) Clustering heat map showing the N-glycoproteins with altered abundances in A391T cortex. 10 mice are shown in the columns, with the first column representing the one TT outlier (*), followed by 5 CC cortex samples and the remaining 4 TT cortex samples. B) PCA analysis of showing that N-glycoproteins from TT mice (black dots) display a clear group difference in cortex compared to controls (white dots). C) Volcano plot of differentially N-glycosylated proteins in A391T cortex. Increased (red) and decreased (blue) N-glycoproteins are shown, with the names of the top 30 included. Significance thresholds for fold change (log2, 2-fold change) and adjusted p-value (−log10) are shown as grey dotted lines on the on the x- and y-axis, respectively. N = 5 mice per genotype.

### Differentially N-glycosylated proteins are enriched in unique cellular components and cell clusters of origin, including glycoproteins implicated in schizophrenia

For further insight into the molecular changes of A391T cortex, we analyzed the list of differentially N-glycosylated proteins using the FUMA GENE2FUNC platform^48^. A list of genes from all detected N-glycoproteins in control cortex (210 genes) and those unchanged (141 gene), decreased (41 genes), and increased (25 genes) in A391T cortex were used as input compared to protein coding genes, all of which were enriched in brain tissue as expected. Gene Ontology (GO) cellular component analysis, based on the PANTHER classification system^49^, again confirmed significant enrichment of predictable components in all lists including dendrite, synapse, and neuron, known locations of N-glycoproteins in the brain. N-glycoproteins decreased in A391T cortex showed a unique enrichment of several components of the plasma membrane and genes involved in ion transport (**Fig S8A**), while N-glycoproteins increased in A391T cortex included multiple components of the ER and Golgi networks (**Fig S8B**). Though there was overlap and the significance of enrichment decreased with the size of the list, these findings suggest the A391T variant has different effects on glycosylation of individual proteins within the same tissue.

Single-cell profiling provides detailed information on the cellular specificity of gene expression in the mouse brain^28^. *Slc39a8* is expressed at relatively low levels in the mouse brain, with expression concentrated in endothelial cells, trace levels in microglia and fibroblasts, and a near total absence in other cell types (**Fig S9A, S9B**). To determine the cell types of origin for differentially N-glycosylated proteins in A391T cortex, we downloaded single cell expression data from mouse frontal cortex using the DropViz platform^28^ for the lists of all, unchanged, decreased, and increased glycoproteins. Total transcript levels were summed within each of the 14 unique cell-type clusters; these data indicated that differentially glycosylated proteins were expressed in all subtypes, with a similar overall pattern observed between all detected glycoproteins in control mice and those unchanged in TT cortex (**Fig S9C**). Normalization of transcript levels for comparison across lists highlighted several differences in the cell cluster of origin, including an enrichment of glycoproteins with decreased levels in interneurons and neurons, and an enrichment of glycoproteins with increased levels in endothelial cells (**Fig 5**), where the majority of *Slc39a8* is expressed.

**Fig. 5.**
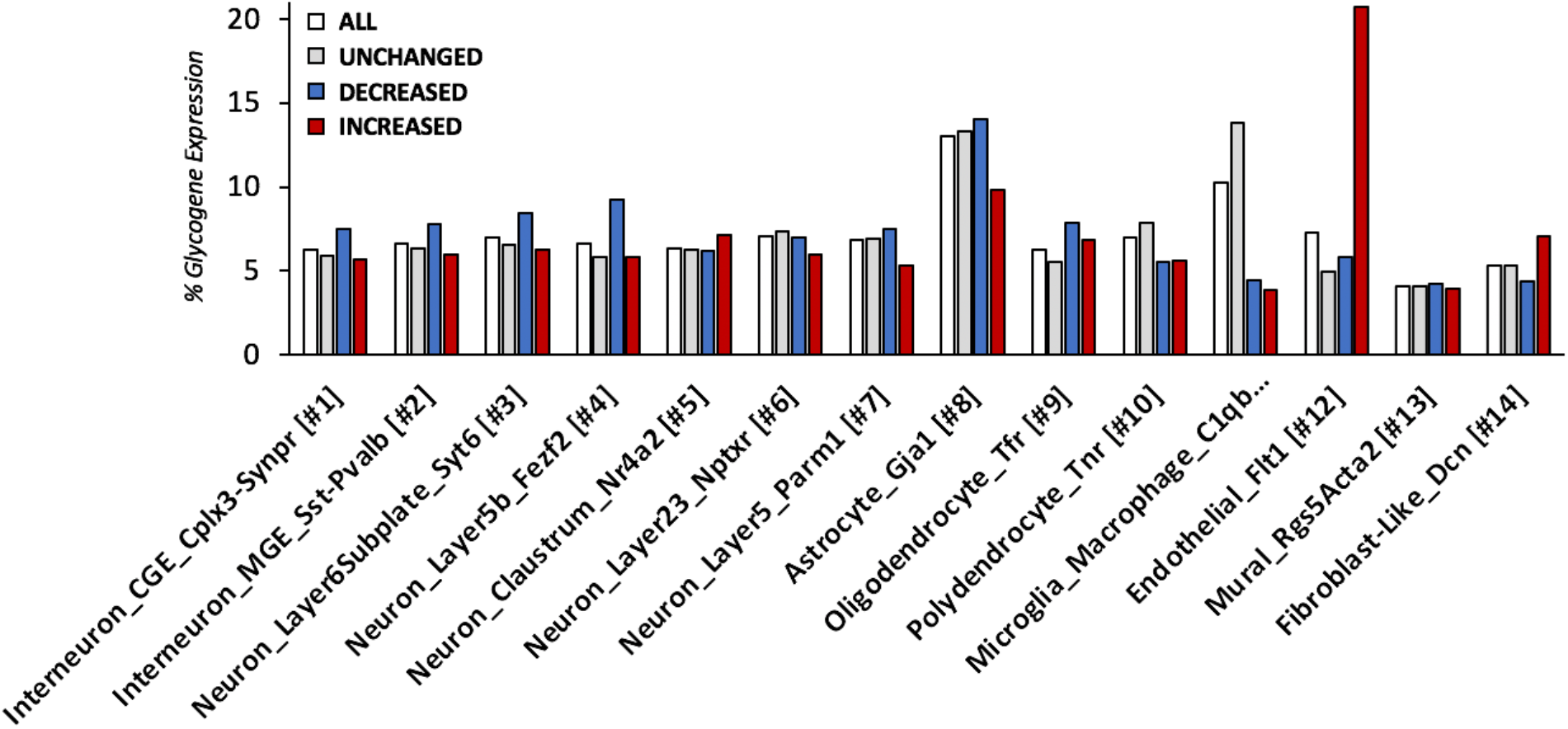
Decreased and increased N-glycoproteins in A391T cortex originate are expressed across cell types in the brain. Single-cell mouse brain expression data for genes encoding differentially N-glycosylated proteins in TT mice were downloaded from www.dropviz.org and compared to all detected N-glycoproteins. After normalization, the sum of clustered transcripts per 100,000 for each distinct cell type is illustrated between groups including all detected glycoproteins (white), unchanged (grey), decreased (blue), and increased (red). Differentially N-glycosylated proteins are expressed across all cell types despite the restricted expression pattern of Slc39a8 in endothelial cells.

Among the 66 differentially glycosylated proteins, several are striking given their function in pathways previously implicated in schizophrenia, including glutamate and GABA neurotransmission (Gria1, Slc1a2, Gabbr1), dopamine signaling (Drd1), and cell adhesion/migration (Cdh5, Ncam1, Ncan, Pcdh19, Reln) (**Table 2**). In addition, Brinp2 and Gria1 were differentially N-glycosylated in A391T cortex and are encoded by genes associated with schizophrenia through GWAS^12^.

**Table 2.** Select N-glycoproteins linked to schizophrenia pathogenesis with altered levels in A391T cortex. A subset of glycoproteins with altered N-glycosylation in the A391T cortex are highlighted, including cell adhesion molecules, proteins involved in neurotransmitter, and two genes associated with schizophrenia by GWAS*. FC, Fold Change; Adj.p.val, Adjusted p-value.

| Gene | Protein | FC | Adj.p.val |
| --- | --- | --- | --- |
| Slc1a2 | Excitatory amino acid transporter 2 | -4.90 | 0.004 |
| Ncam1 | Neural cell adhesion molecule 1 | -2.51 | 0.007 |
| Reln | Reelin | -2.42 | 0.006 |
| Gabbr1 | Gamma-aminobutyric acid type B receptor subunit 1 | -1.97 | 0.017 |
| Cdh5 | Cadherin-5 | -1.89 | 0.004 |
| Pcdh19 | Protocadherin-19 | -1.58 | 0.044 |
| Gria1* | Glutamate receptor 1 | 1.61 | 0.002 |
| Ncan | Neurocan core protein | 1.86 | 0.025 |
| Brinp2* | BMP/retinoic acid-inducible neural-specific protein 2 | 2.10 | 0.013 |
| Drd1 | D(1A) dopamine receptor | 2.15 | 0.002 |

## Discussion

Here we employed several sensitive techniques to demonstrate that in a murine model the schizophrenia risk allele in *SLC39A8* reduces the products of manganese-dependent Golgi-glycosyltransferases in the brain, with a relatively small effect size as predicted for common genetic variants. Rodent studies are not a model for schizophrenia but can be a tool to understand downstream effects of risk variants. In contrast to previous studies which have primarily knocked out schizophrenia risk genes in mice, these results show that the biologically relevant risk allele in *SLC39A8* alters key molecular pathways in the brain.

Protein glycosylation changes in A391T mice were most robust in the N-glycosylation pathway, which contains several Mn^2+^-binding glycosyltransferases with DxD motifs, such as β-1,4-galactosyltransferases and N-acetylglucosaminyltransferases^19,20^. In addition, oligosaccharyltransferase (OST), which transfers dolichol anchored N-glycan precursors to asparagine residues of nascently transcribed proteins in the ER lumen, also requires Mn^2+^ for activity^50,51^. Though there was an increase in the relative abundance of tri- and tetra-antennary N-glycans in the cortex of A391T mice by MALDI, quantitative studies using F-MAPA confirmed a reduction of total N-glycans, and suggest the cortex prioritizes synthesis of these low abundance structures when pathway activity is reduced. A391T mice also exhibited small changes in brain O-glycosylation, and the twenty N-acetylgalatosaminyltransferases (*GALNTs*), which initiate O-GalNAc-type glycosylation, contain a Mn^2+^-binding DxH domain and could be affected by changes in Mn^2+^ levels^52^. Transcriptomics from cortex and cerebellum did not identify changes that could account for the observed glycosylation differences in A391T brain, as the RNA levels of glycosyltransferases, glycosylhydrolases, and the differentially glycosylated targets were unchanged. This observation is consistent with human data as the rs13107325 SNP is not an eQTL for any gene in the brain including *SLC39A8*^26^.

Glycosylation changes associated with the A391T variant support the converging evidence that dysglycosylation is involved in schizophrenia pathogenesis. Numerous glycosylation enzymes are associated with schizophrenia through GWAS^12,18^ and the number continues to expand, with many glycosylation genes included in the most recent list of prioritized schizophrenia risk genes^13^, including several specific to N-glycosylation (*MAN2A1*, *ALG12*, *MANBA*) and O-glycosylation (*GALNT10*, *GALNT17*/*WBSCR17*, *TMTC1*). In addition, many genes associated with schizophrenia through genetic studies encode proteins that are critically regulated by glycosylation^18^, and post-mortem studies have consistently shown brain dysglycosylation in individuals with schizophrenia^53^. Glycoproteomic changes further illustrate the connection between schizophrenia and glycosylation, as several risk genes and proteins involved in critical developmental and functional pathways in the brain are altered in the cortex of A391T mice. Pathway analyses of the differentially glycosylated proteins suggest that proteins with increased or decreased levels are enriched in distinct pathways and cellular components, consistent with the complexity of glycosylation as a non-template-based protein modification. We predict glycosylation enzymes exert altered substrate specificity and binding in the setting of homeostatic changes in Mn^2+^ levels, increasing the glycosylation of some priority targets while decreasing others.

*SLC39A8* is expressed in most tissues at low levels^27^, and in the mouse brain *Slc39a8* is restricted primarily to endothelial cells^28^. We observed changes in glycoproteins originating from, though not necessarily restricted to, all cell clusters in the brain. Interestingly, glycoproteins whose levels are increased in TT cortex are enriched in endothelial cells, which express the highest level of *Slc39a8*, while decreased glycoproteins are enriched in several neuronal clusters, with a minimal change in microglia. These results suggest that Slc39a8 serves as a gate keeper for Mn^2+^ in the brain, with the A391T variant impairing its function in endothelial cells to result in subtle but significant glycosylation changes across the brain and cell types. It is possible that the biological effects are related to its peripheral expression, but the final consequence is altered brain glycosylation.

We confirmed our prior observation of a sex-dependent effect of A391T on glycosylation, with human and mouse males showing a larger effect compared to females^42^. Recent investigations of gastrointestinal phenotypes associated with A391T confirmed sex effects, which may in part be explained by lower Mn^2+^ levels in male mice^43,54^. It is important to note that the increased risk for schizophrenia in A391T carriers is equal in males and females^13^. We suspect that A391T confers risk through altered Mn^2+^ transport during brain development; given the complexity of Mn^2+^ homeostasis and transport, the subtle effect may normalize over time despite vulnerable pathways during critical windows having been already affected. Interestingly, we did not observe glycosylation changes in male A391T mice at 4 weeks of age, which roughly corresponds with early adolescence in humans, raising the possibility that Mn^2+^ supplementation after 4 weeks of age might prevent these changes.

In sum, our findings provide mechanistic support that the *SLC39A8* schizophrenia risk allele alters glycosylation in the brain and provide several nodes of connection between known genetic risk and disease pathways. Mendelian disorders of glycosylation caused by severe hypofunctioning alleles of SLC39A8 are treated with supplementation of glycosylation precursors and manganese^22,23^. Although the A391T variant is likely to have a much smaller effect on SLC39A8 function, supplementation with similar factors during critical periods of brain development may lessen the risk of neuropsychiatric conditions in carriers and targeting glycosylation more broadly in schizophrenia could represent a novel pathway for therapeutic development.

## Author Contributions

RGM conceptualized the project, performed the bulk of experiments and analyses, and wrote the manuscript

SEW performed glycomics experiments and assisted statistical analysis

MN performed quantitative glycan analyses and lectin blotting

BY and ADS performed glycoproteomic analysis

TN assisted in tissue harvest, genotyping and mouse colony maintenance

DBG was involved in mouse generation

EAC was involved in mouse generation, genotyping and colony maintenance

MC performed analysis of RNAseq data

RS supervised RNAseq analysis

EMS initiated the project and coordinated collaborations

CMW supervised BY and ADS and oversaw glycoproteomic analysis

JWS co-supervised RGM and SEW, oversaw experimental analyses, and helped conceptualize the project

RJX provided mouse samples for analysis

RDC co-supervised RGM, SEW and MN, oversaw experimental analyses, and helped conceptualize the project

All authors contributed feedback and edits to the manuscript

## Acknowledgements

This work was supported by a foundation grant from the Stanley Center for Psychiatric Research at the Broad Institute of Harvard/MIT (awarded to RGM) and NIH grant P41GM103694 (awarded to RDC). RGM is supported by T32MH112485. CMW is supported by NIH NCI U01CA242098.

## Methods

*A391T homozygous knock-in mutant mice* were generated by gene targeting in embryonic stem cells derived from C57BL/6J mice, as previously described^43^. Mice from both sexes were used in this study and were 12 weeks old at the time of tissue harvest unless otherwise noted. No live animals were used in this study. All mice were housed and maintained in accordance with the guidelines established by the Animal Care and Use Committee at Massachusetts General Hospital under protocol #2003N000158.

*Sample Preparation and Glycomic Analysis* were performed as previously described in detail in a related manuscript^44^.

### F-MAPA derivation and quantification of released N-glycans

All chemical reagents were HPLC grade purchased from Sigma Aldrich (St. Louis, MO, USA) unless otherwise noted, and all reagents were freshly prepared on the day of analysis. PNGase F (New England Biolabs, #P0704) released N-glycans from 1 mg of brain protein lysate were isolated as previously described^44^. Purified N-glycans were derivatized using the fluorescent linkerfluorenylmethyloxycarbonate-3-(methoxyamino)propylamine (F-MAPA) as previously described^46^. In brief, purified N-glycans (not permethylated) were resuspended in 80 μL of dimethylsulfoxide/acetic acid at 7:3 (v/v) ratio containing 0.25 M F-MAPA, 0.25 M sodium acetate, and 0.5 mM 2-amino-5-methoxybenzoic acid and incubated for 2 hours at 65°C with gentle rotation at 100 RPM. Ten (10) volumes of ethyl acetate were added, vortexed, and incubated at −20°C for 20 minutes to precipitate N-glycan/F-MAPA conjugates and isolated by centrifugation at 10,000 RPM for 15 minutes. After removal of the supernatant, the pellet containing F-MAPA derived N-glycans was dissolved in 300 μL of Milli-Q filtered water and loaded on a C18 Sep-Pak columns (200 mg) preconditioned with acetonitrile (ACN) then water. Bound F-MAPA-linked N-glycans were washed with 5 mL of water, eluted with 2 mL of 30% ACN, and lyophilized. Purified F-MAPA linked N-glycans were then resuspended in 200 μL of water and detected using a fluorescence spectrophotometer (SpectraMax i3x, Molecular Devices) with excitation wavelength of 265 nm and emission wavelength 315 nm and quantified (SoftMax Pro Software, v7.0.3).

### Sialic acid quantification

Absolute quantification of sialic acid in different glycoconjugate fractions was performed using the Sialic Acid (NANA) Assay Kit (Ab83375, Abcam, Cambridge, MA) according to the manufacturer’s instructions. Briefly, 30 μg of protein or 600 pmol of N-glycan/F-MAPA were incubated with the sialidase NeuA for 4 h at 37°C. The reaction was stopped by boiling the samples for 5 minutes at 95°C, followed by centrifugation at 10,000 RPM for 5 minutes, and the supernatant (containing free sialic acid) was transferred to a 96-wells plate. NANA reactions were performed by adding 25 μL reaction mix (22 μL assay buffer, 1 μL sialic acid converting enzyme, 1 μL sialic acid development mix, 1 μL sialic acid probe) to each supernatant, and incubated for 30 minutes at room temperature in the dark. The fluorescence of each well was then quantified by spectrophotometry (Excitation: 535 nm, Emission: 587 nm) as described above.

*RNA Sequencing* was performed as described performed as previously described in related manuscript^44^. In brief, RNA from snap frozen cortex and cerebellum was purified using the RNeasy Lipid Tissue Mini Kit (QIAGEN, 74804). RNA-seq libraries were prepared from total RNA using polyA selection followed by the NEBNext Ultra II Directional RNA Library Prep Kit protocol (New England Biolabs, E7760S). Sequencing was performed on Illumina HiSeq 2500 instrument resulting in approximately 30 million of 50 bp reads per sample. Sequencing reads were mapped in a splice-aware fashion to the mouse reference transcriptome (mm9 assembly) using STAR ^55^. Read counts over transcripts were calculated using HTSeq based on the Ensembl annotation for GRCm37/mm9 assembly and presented as Transcripts Per Million (TPM) ^56^.

### N-Glycoproteomic analysis

Brain cortex samples were lysed in 1 mL lysis buffer (20 mM HEPES pH 7.9, 1% SDS, and protease inhibitor) with protein concentrations determined by BCA assay (Thermo Fisher Scientific, Cat # 23225). Reduction and alkylation were performed as previously described50. S-trap digestion was performed according to the manufacturer’s instructions (ProtiFi, Cat # C02-mini), resulting in ~0.6 mg tryptic peptides per sample. To one half of tryptic peptides, 15 mg of HILIC beads (PolyLC, Cat # BMHY0502) pre-activated with 0.1% trifluoroacetic acid (TFA) were added to make a 1:50 peptide-to-beads mass ratio. The samples were vortexed in binding buffer (0.1% TFA, 19.9% water, 80% ACN) for 1 h at room temperature to allow N-glycopeptides to bind to beads. The unbound peptides were washed with 150 μL binding buffer 6 times, and N-glycopeptides were eluted by washing the beads with (150 μL) 0.1% TFA for 5 times. Finally, 2 μL PNGase F (500U/μL) (NEB # P0704) was added to the elution buffer and the samples were incubated for 3 h at 37 °C, followed by lyophilization. To the other half of tryptic peptides, multi-lectin enrichment was performed based on a previously reported protocol with some modifications51. Briefly, tryptic peptides were mixed with a mixture of lectins (90 μg ConA, 90 μg WGA, 36 μg RCA120 in 2X binding buffer (Sigma-Aldrich, Cat L7886, Cat L9640, and Cat L7647) and transferred to a 30 kDa filter (Pall Corporation, Cat # OD030C34), incubated at room temperature for 1 h and unbound peptides were eluted by centrifuging at 14,000 g for 10 minutes. N-glycopeptides were washed with 200 μL binding solution four times and 50 μL digest buffer of 50 mM triethylammonium bicarbonate (TAEB) twice. Finally, 2 μL PNGase F (NEB # P0704) was added to the filter and incubated for 3 h at 37 °C. The deglycosylated N-glycopeptides were eluted with 2 × 50 μL digest buffer. N-glycopeptides from two enrichment methods were combined and desalted by C18 tips (Thermo Fisher Scientific, Cat #87784) following the manufacturer’s instructions and resuspended in 30 μL 100 mM TEAB buffer. For each sample, 5 μL the corresponding amine-based tandem mass tag (TMT) 10-plex reagents (10 μg/μL) (Thermo Fisher Scientific, Cat # 90406) was added and incubated for 1 hour at room temperature. The reactions were quenched with 2 μL 5% hydroxylamine solution (Oakwood Chemical, Cat # 069272), and the combined mixture was lyophilized. High-pH fractionation was done according to the manufacturer’s instructions (Thermo Fisher Scientific, Cat # 84868), which resulted in 15 independent fractions.

A Thermo Scientific EASY-nLC 1000 system was coupled to a Thermo Scientific Orbitrap Fusion Tribrid with a nano-electrospray ion source. Mobile phases A and B were water with 0.1% formic acid (v/v) and ACN with 0.1% formic acid (v/v), respectively. For each fraction, peptides were separated with a linear gradient from 4 to 32% B within 45 min, followed by an increase to 50% B within 10 minutes and further to 98% B within 10 minutes, and re-equilibration. The instrument parameters were set as follows: survey scans of peptide precursors were performed at 120K FWHM resolution over a m/z range of 410-1800. HCD fragmentation was performed on the top 10 most abundant precursors exhibiting a charge state from 2 to 5 at a resolving power setting of 50K and fragmentation energy of 37% in the Orbitrap. CID fragmentation was applied with 35% collision energy and resulting fragments detected using the normal scan rate in the ion trap.

The raw data was processed using Proteome Discoverer 2.4 (Thermo Fisher Scientific). Data was searched against the UniProt/SwissProt mouse (Mus musculus) protein database (May 30, 2019, 17,372 total entries) and contaminant proteins using Sequest HT and Byonic algorithms. Searches were performed with the following guidelines: spectra with a signal-to-noise ratio greater than 1.5; trypsin as enzyme, 2 missed cleavages; variable oxidation on methionine residues (15.995 Da) and deamidation on asparagine (0.984 Da); static carboxyamidomethylation of cysteine residues (57.021 Da), static TMT labeling (229.163 Da) at lysine residues and peptide N-termini; 10 ppm mass error tolerance on precursor ions, and 0.02 Da mass error on fragment ions. Data were filtered with a peptide-to-spectrum match (PSM) of 1% FDR using Percolator. The TMT reporter ions were quantified using the Reporter Ions Quantifier with total peptide normalization. For the obtained PSMs, the data was further filtered with the following guidelines: confidence is high; PSM ambiguity is unambiguous; modifications contain deamidated; exclude all contaminant proteins. Consensus sequence filtering was applied using a string match module with filtering the Subcellular location for Cell membrane, Endoplasmic reticulum, Golgi apparatus, Extracellular, and Secreted. Most of the empty Abundances, if any, were filled in with minimum noise level computed by taking the minimum for each channel in CC and TT cortex. Reducing these minimum groups, five channels for CC and five for TT, required taking the absolute maximum in CC and TT and the absolute minimum in CC and TT. Subsequently, 2000 centroids were generated at random from these 4 points and a minimum noise level was generated using a K-means clustering method. If there were few missing values the maximum and minimum of the Abundances for that PSM in the CC and TT groups were used to generate centroid data. If only one Abundance was missing, the instance was filled with the geometric mean of the PSM for CC or TT. If all Abundances were missing for CC and TT, the PSM was removed. Based on the algorithm in this script of an empty abundance missing completely at random, missing not at random, or missing at random, any valid instances were filled with the appropriate method described above. A limitation of the log-ratio of raw MS data is the dependence on variation within each channel for CC and TT. Applied here was a VSN normalization computed on the imputed matrix using a robust variant of the maximum-likelihood estimator for an additive-multiplicative error model and affine calibration. The model incorporates dependence of the variance on the mean intensity and a variance stabilizing data transformation. The MS data were deposited at the ProteomeXchange Consortium52 via the PRIDE partner repository and are available with the identifier PXD021632.

*Glycoprotein pathway analyses* was performed using the GENE2FUNC tool of the FUMA platform^48^ on October 30^th^, 2020. Input lists included all detected N-glycoproteins in CC cortex, unchanged N-glycoproteins in TT cortex, decreased N-glycoproteins in TT cortex, and increased N-glycoproteins in TT cortex, based on corrected p-values < 0.05 and fold change >2.0 compared to protein coding genes as background. Output results for gene set enrichment of GO cellular components^49^ were presented.

*Glycoprotein cell of origin analyses* was performed using publicly available data downloaded from the DropViz database^28^ on October 30^th^, 2020. Single cell expression data from each gene in the above-described lists of N-glycoproteins were downloaded from frontal cortex containing 14 distinct cell clusters and exponentially natural log transformed to approximate transcripts per 100,000 in each cluster. Transcripts levels for each cluster were then summed for the above described lists of detected glycoproteins (*all, unchanged, decreased, increased*) and normalized within each group to generate the relative abundance of glycogene expression from each cluster.

### Statistical Analysis

For glycomic studies, the abundance of individual glycans and glycan classes were compared between CC and TT mice using unpaired t-tests assuming unequal variance. For RNAseq results, the EdgeR method was used for differential expression analysis with gene cutoffs of 2-fold change in expression value and false discovery rates (FDR) below 0.05 as previously described^57^. For glycoproteomics results, a linear model was fitted to the expression data for CC and TT cortex, and t-statistics were computed by empirical Bayes moderation of standard errors towards a common value. The heatmap clustering method used was based on average linkage.

