## Supplemental Figures for "The schizophrenia-associated variant in *SLC39A8* alters N-glycosylation in the mouse brain"

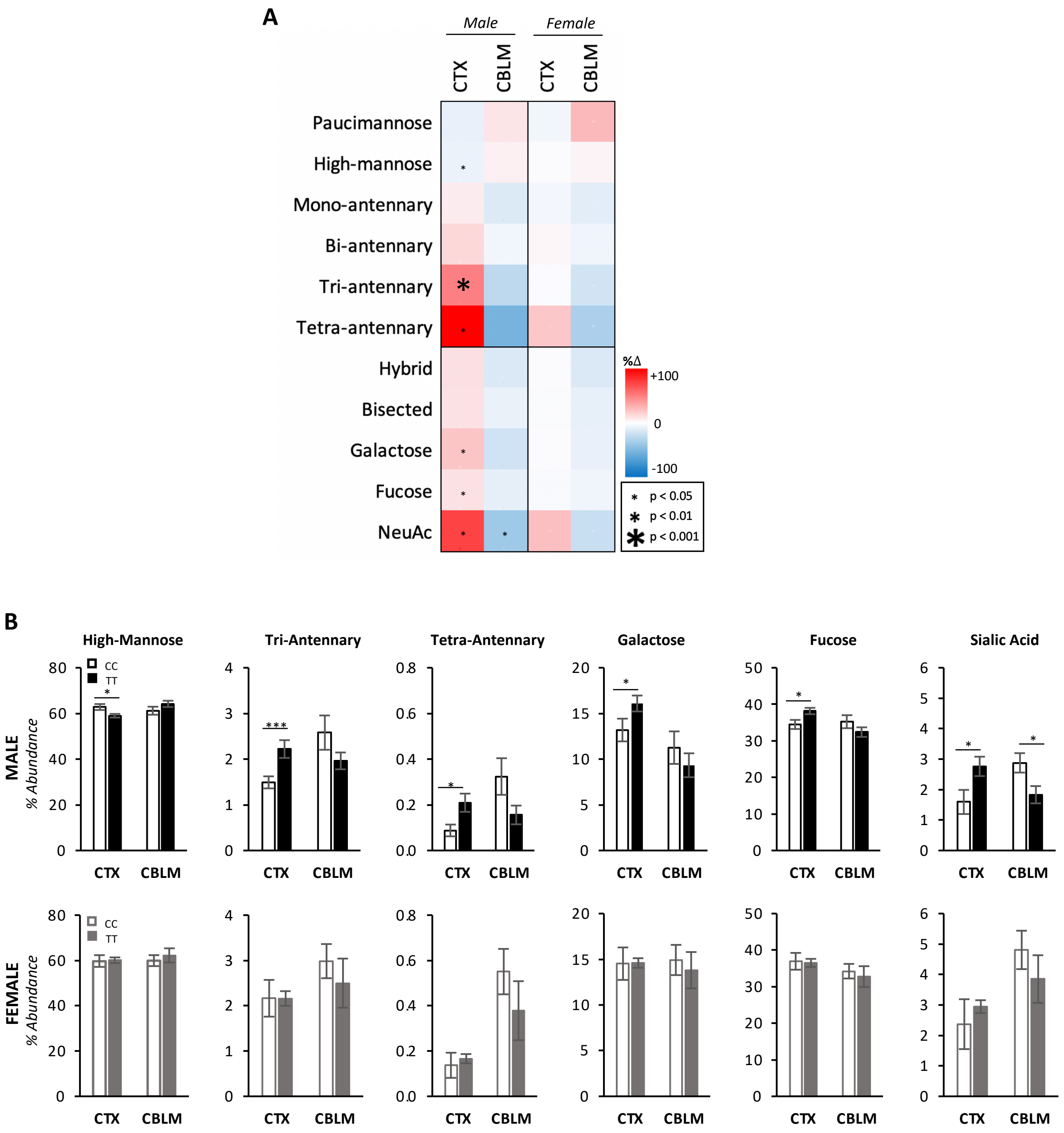


**Fig. S1. A391T shows a stronger effect on brain N-glycosylation in males.** A) Categorical analysis of CTX and CBLM N-glycans from female A391T mice showed the same directionality of change as observed in males, but this effect does not reach statistical significance. B) A391T mice showed altered abundance of multiple N-glycan categories. Data presented as a heat map of percent change in glycan abundance comparing TT mice to CC controls. For N-glycans, CC=4, TT=4 for each sex. Data presented as mean percent abundance +/- SEM. Related to Fig. 1.

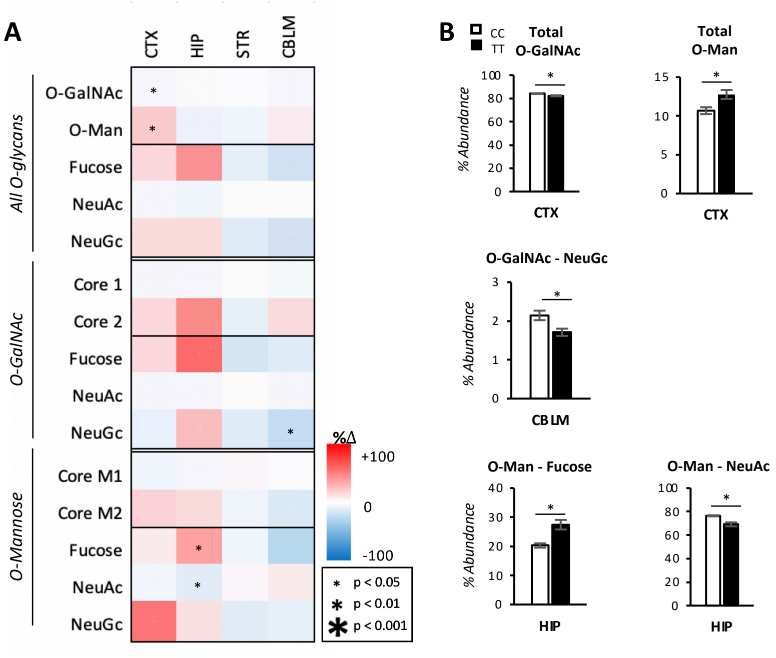


**Fig. S2. A391T has a smaller effect on the relative abundance of protein O-glycans.** A) Categorical analysis of O-glycans from four brain regions of male mice revealed region-specific changes in certain glycan classes. Data presented as a heat map of percent change in glycan abundance comparing TT mice to CC controls. B) Cortex, hippocampus, and cerebellum from A391T mice showed altered abundance of certain O-glycan categories. CTX CC=4, TT=5; HIP CC=2, TT=4; STR CC=4, TT=6; CBLM CC=4, TT=5. Data presented as mean percent abundance +/- SEM. Related to Fig. 1.

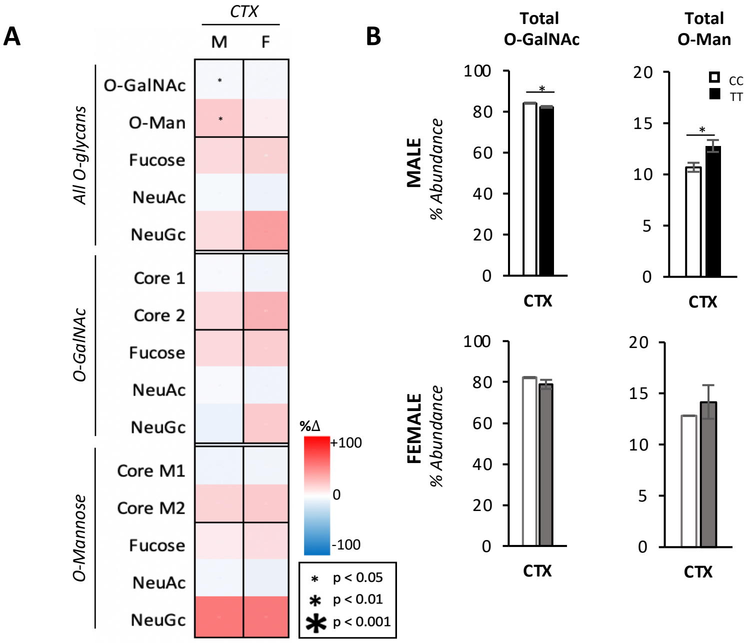


**Fig. S3. A391T shows a stronger effect on O-glycosylation in males.** A) Categorical analysis of CTX and CBLM O-glycans from female A391T mice showed the same directionality of change as observed in males, but this effect does not reach statistical significance. Data presented as a heat map of percent change in glycan abundance comparing TT mice to CC controls. N = 2 CC, 4 TT for each region and gender. Data presented as mean percent abundance +/- SEM. Related to Fig. 1.

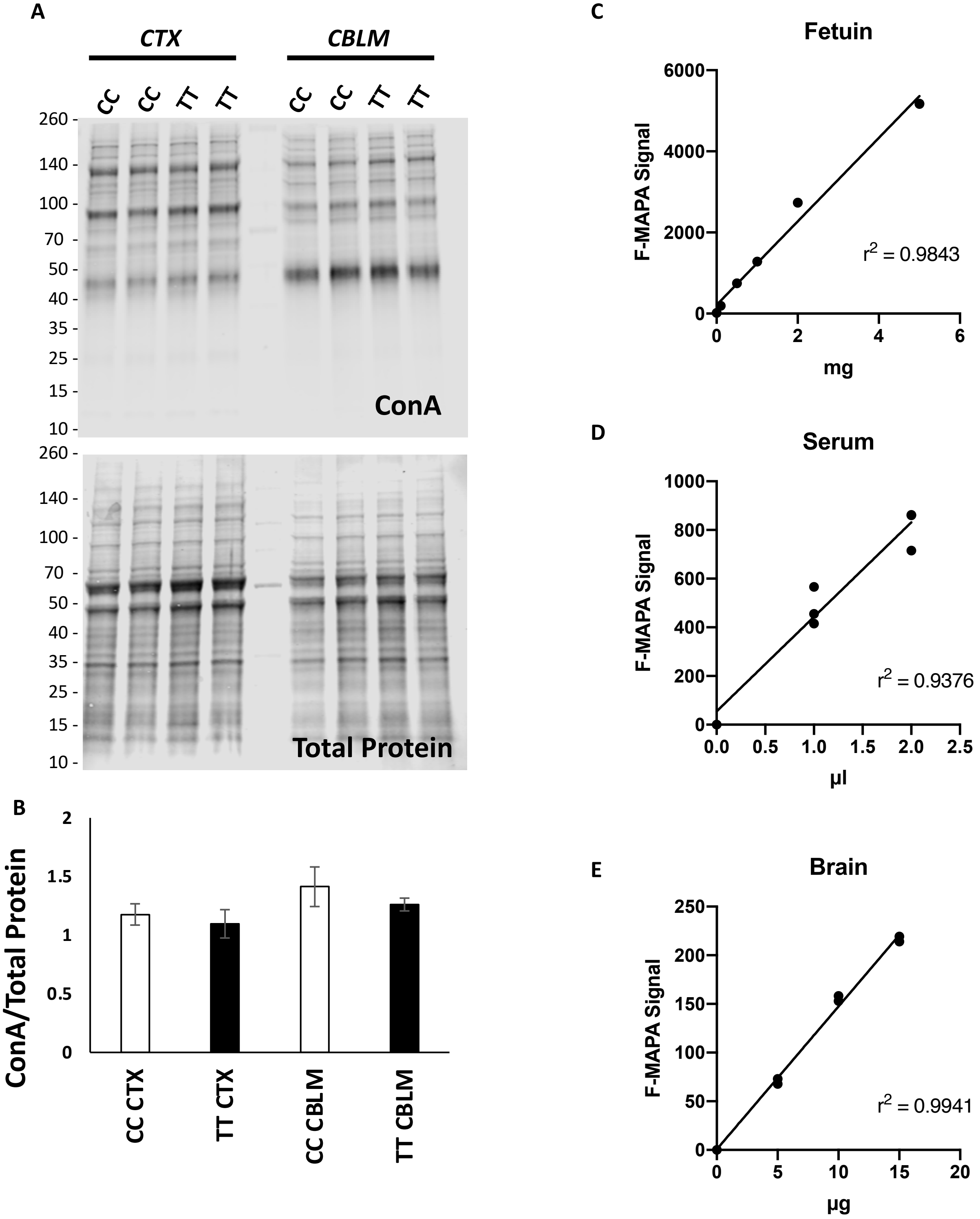


**Fig. S4. Quantitative measures of brain glycans.** A) Representative lectin western blot analysis of brain lysate using ConA and total protein stain from cortex and cerebellum of male CC and TT mice. B) Quantification of ConA/Total Protein signal from western blots illustrated above. N = 4 per region and genotype. Data shown as mean +/- SEM. F-MAPA fluorescence curves using different quantities or concentrations of standards controls of fetuin (C), serum (D), and brain lysate (E), with individual data points and the corresponding $r^2$ values. Related to Fig. 2.

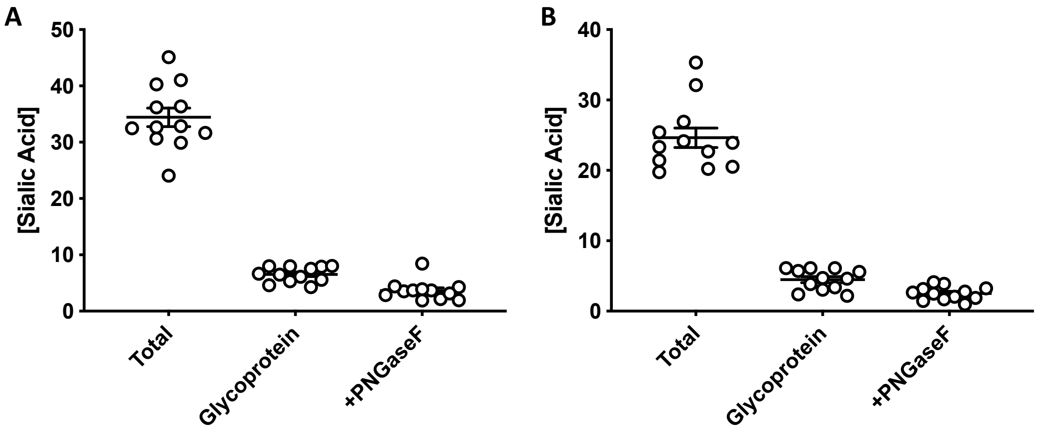


**Fig. S5. Most brain sialic acid is contained on glycolipids.** The concentration of sialic acid in cortex (A) and cerebellum (B) of wild-type mice, presented as nmol sialic acid/mg of protein, determined using the NANA kit (Abcam) and BCA assay (Pierce). Fractions represents crude brain homogenate (Total), N- and O- glycoproteins following glycolipid extraction with methanol/chloroform (Glycoprotein), and O-glycoproteins after PNGase F treatment (+PNGaseF). Individual data points are shown, with brackets representing group means +/- SEM. Related to Fig. 2.

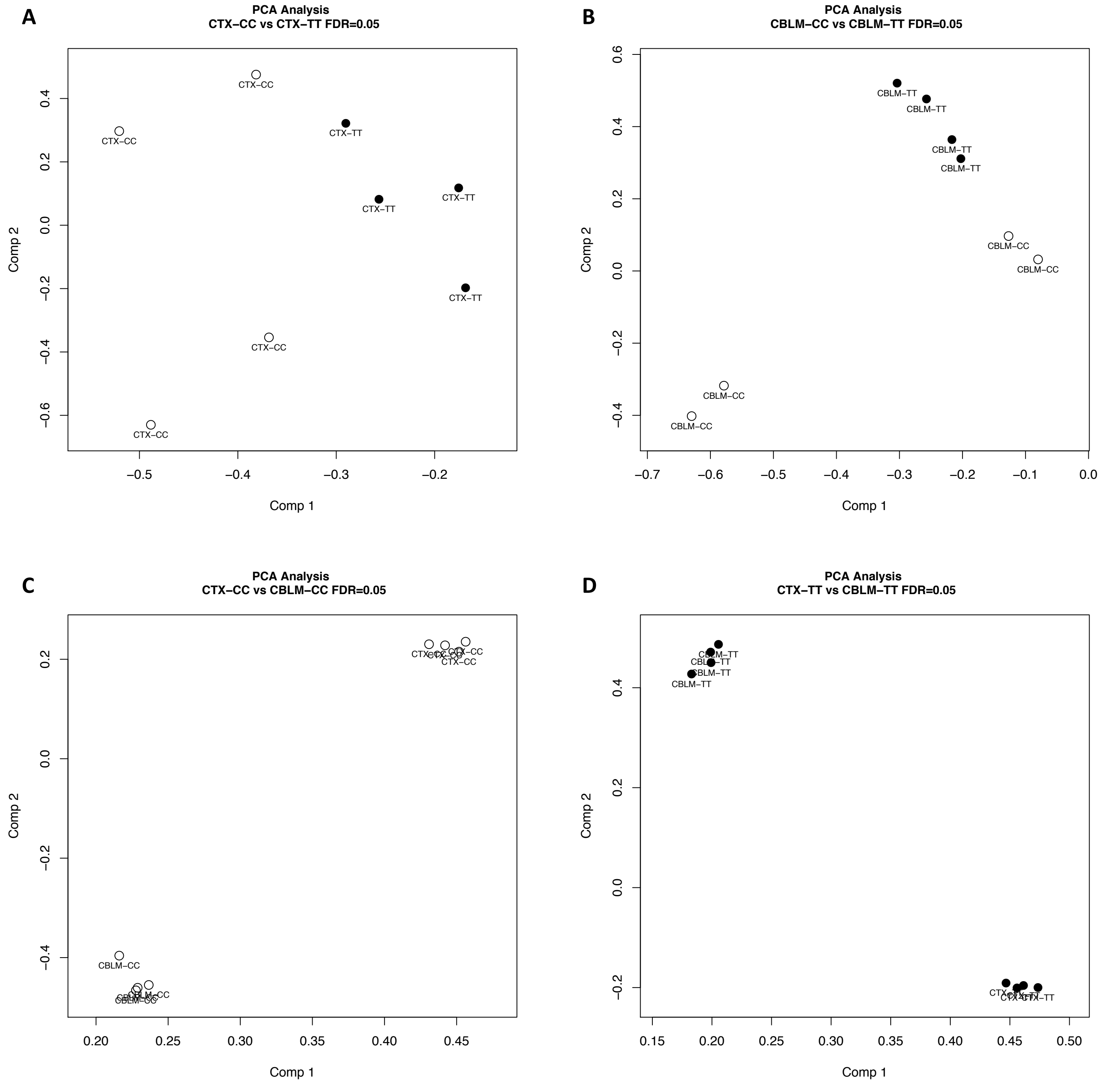


**Fig. S6. Gene expression changes are minimal between genotypes but large between regions.** PCA plots comparing cortex (A) and cerebellum (B) between CC and TT mice identifies minimal gene expression changes between genotypes. PCA plots of CC (C) and TT (D) mice between cortex and cerebellum shows large gene expression changes, consistent with the known regional differences. N = 4 male mice per group. Related to Fig. 3.

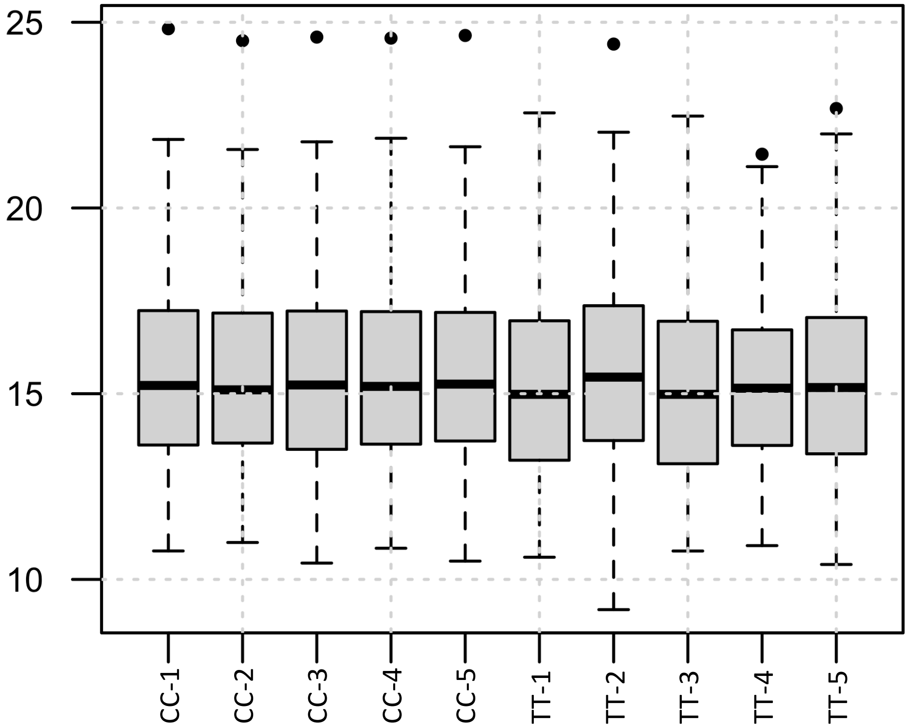


**Fig. S7. Normalization Boxplots for N-glycoproteomic analysis.** The distribution of abundances for each channel incorporating dependence of the variance on the mean intensity and a variance stabilizing data transformation to account for any variation in technical replicates. Overall, the Control (CC) and Treatment (TT) has a similar distribution across replicates validating the statistical significance of the given glycoproteomics results, with only the known outlier TT-2 showing slightly increased variance compared to other samples.

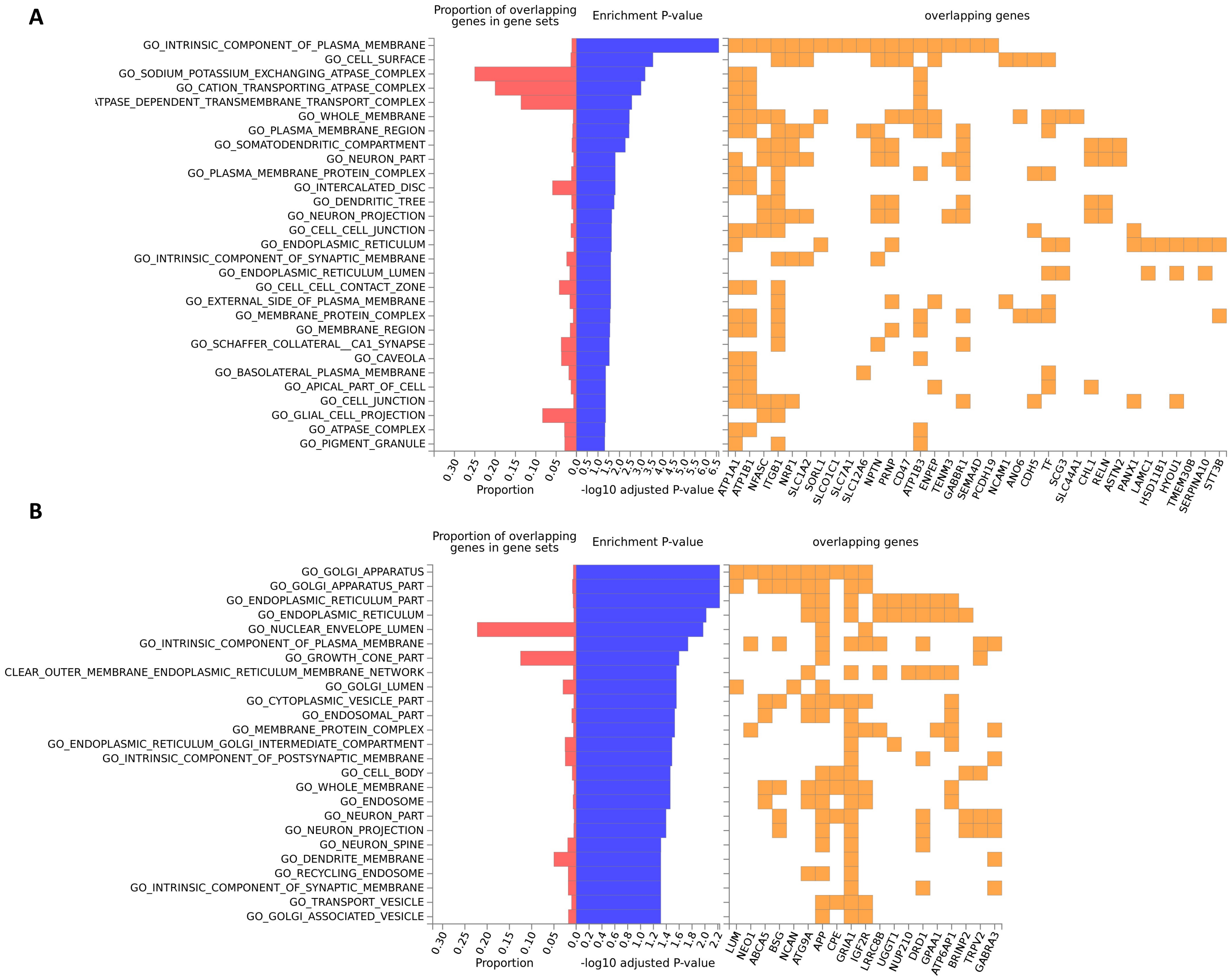


**Fig. S8. Decreased and increased N-glycoproteins in A391T cortex are enriched in unique cellular components.** A) GO analysis of N-glycoproteins decreased in A391T cortex shows an enrichment of proteins in the plasma membrane and involved in ion transport. B) GO analysis N-glycoproteins increased in A391T cortex shows an enrichment of proteins in the ER and Golgi apparatus. Adjusted p-values <0.05 (-log10) were calculated using the FUMA GENE2FUNC platform.

A

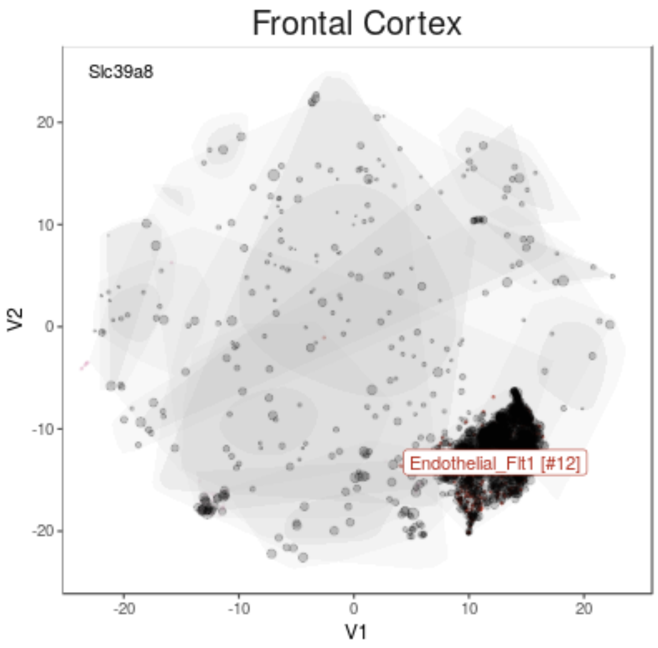


B

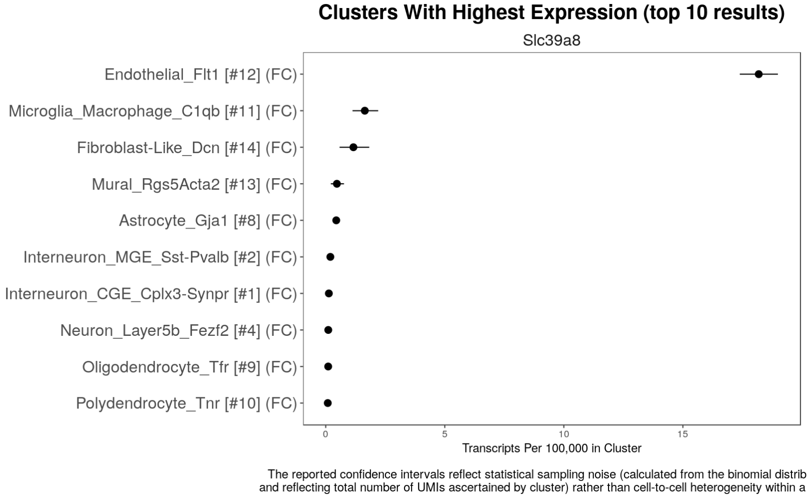


C

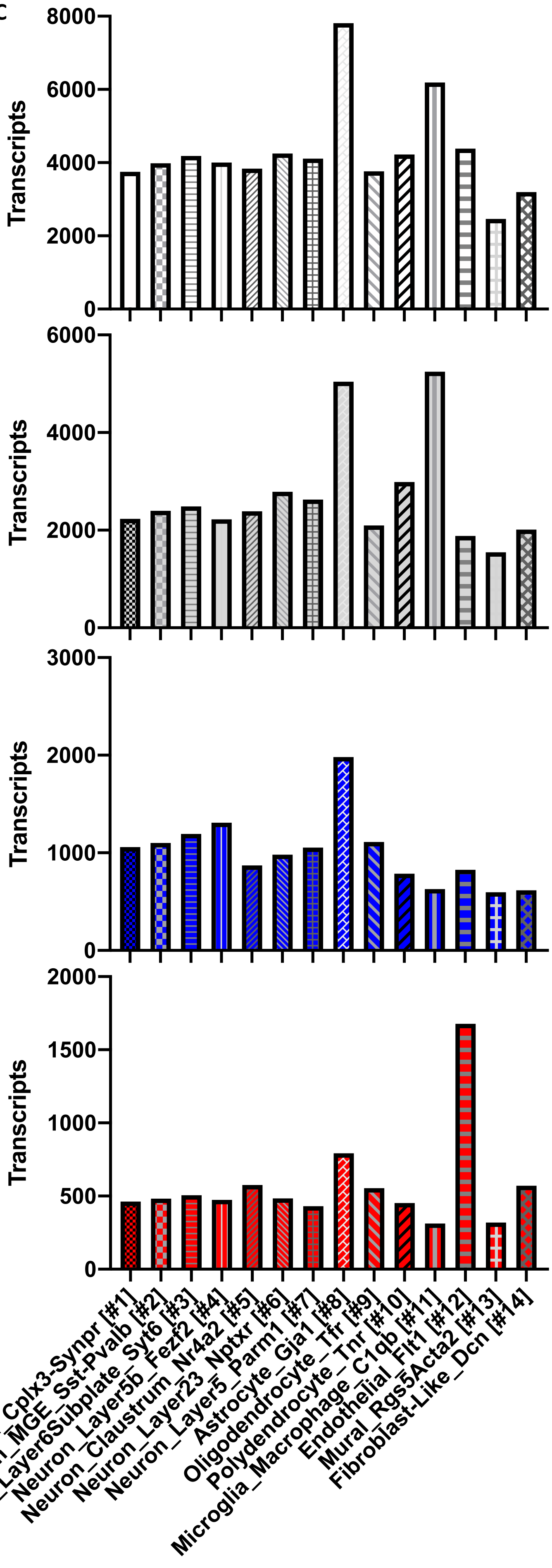


**Fig. S9.** ***Slc39a8*** **is enriched in endothelial cells, though N-glycoproteins across all cell clusters are affected by A391T.** A) tSNE analysis of Slc39a8 in frontal cortex of mice shows dramatic enrichment in enrichment in endothelial cells. B) Cluster analysis of Slc39a8 mouse cortex shows the high level of RNA expression in endothelial cells, with trace levels in microglia and fibroblasts, and a near total absence of expression in other cell types including neurons, astrocytes, and oligodendrocytes. C) Single-cell mouse brain expression data for genes encoding differentially N-glycosylated proteins in TT mice are represented across all cell clustering. Data for glycoprotein expression was downloaded from www.dropviz.org, with the sum of Clustered transcripts per 100,000 for all detected glycoproteins (white), unchanged (grey), decreased (blue), and increased (red) within each distinct cell type.
