## Supplemental Tables for "The schizophrenia-associated variant in *SLC39A8* alters N-glycosylation in the mouse brain"

Mealer, *et al.*

**Supplemental Tables**

Table S1. Comparison of individual N-glycan abundance in CC vs TT brain regions
Table S2. Brain N-glycan structure, name, mass, and characteristics used in study
Table S3. Categorical analysis of protein N-glycans in 4-week old mice shows no difference in CTX and CBLM
Table S4. Brain O-glycan structure, name, mass, and characteristics used in study
Table S5. Comparison of individual O-glycan abundance and categories in CC vs TT brain regions
Table S6. Differentially expressed genes in cortex and cerebellum of CC vs TT mice at FDR of 0.05
Table S7. N-glycoprotein levels in cortex of TT mice relative to CC controls. FC, fold change

**Table S1. Comparison of individual N-glycan abundance in CC vs TT brain regions.**

| | *Male* | | | | | | | | | | | |
|---|---|---|---|---|---|---|---|---|---|---|---|---|
| | **Cortex** | | | **Hippocampus** | | | **Striatum** | | | **Cerebellum** | | |
| ***m/z*** | CC | TT | *p-value* | CC | TT | *p-value* | CC | TT | *p-value* | CC | TT | *p-value* |
| 1141.6 | 0.657 | 0.571 | 0.3 | 0.556 | 0.590 | 0.7 | 0.647 | 0.416 | 0.1 | 0.286 | 0.322 | 0.5 |
| 1171.1 | 0.223 | 0.202 | 0.5 | 0.190 | 0.199 | 0.7 | 0.220 | 0.182 | 0.2 | 0.100 | 0.121 | 0.3 |
| 1345.8 | 1.576 | 1.465 | 0.5 | 1.184 | 1.126 | 0.7 | 1.108 | 1.038 | 0.6 | 0.452 | 0.491 | 0.5 |
| 1375.8 | 0.390 | 0.376 | 0.7 | 0.402 | 0.382 | 0.6 | 0.370 | 0.339 | 0.3 | 0.197 | 0.217 | 0.4 |
| 1416.8 | 0.108 | 0.110 | 0.9 | 0.119 | 0.115 | 0.6 | 0.113 | 0.104 | 0.6 | 0.068 | 0.070 | 0.8 |
| 1549.9 | 0.253 | 0.256 | 0.9 | 0.265 | 0.236 | 0.1 | 0.271 | 0.242 | 0.4 | 0.135 | 0.127 | 0.6 |
| 1579.9 | 46.043 | 42.962 | 0.1 | 46.263 | 42.734 | 0.1 | 45.396 | 46.198 | 0.6 | 37.135 | 41.082 | 0.1 |
| 1590.9 | 0.505 | 0.503 | 0.9 | 0.470 | 0.441 | 0.2 | 0.372 | 0.381 | 0.7 | 0.170 | 0.162 | 0.5 |
| 1620.9 | 0.142 | 0.142 | 1.0 | 0.153 | 0.144 | 0.3 | 0.147 | 0.137 | 0.5 | 0.118 | 0.124 | 0.7 |
| 1661.9 | 0.490 | 0.529 | 0.3 | 0.503 | 0.459 | 0.1 | 0.403 | 0.413 | 0.7 | 0.189 | 0.187 | 0.9 |
| 1754.0 | 0.175 | 0.176 | 0.8 | 0.206 | 0.183 | 0.1 | 0.180 | 0.173 | 0.7 | 0.138 | 0.126 | 0.4 |
| 1784.0 | 7.722 | 7.229 | 0.1 | 6.864 | 6.600 | 0.4 | 7.247 | 7.260 | 1.0 | 13.443 | 13.885 | 0.4 |
| 1795.0 | 0.407 | 0.417 | 0.7 | 0.443 | 0.407 | **0.044*** | 0.429 | 0.435 | 0.8 | 0.236 | 0.220 | 0.3 |
| 1825.0 | 0.314 | 0.320 | 0.8 | 0.344 | 0.318 | 0.1 | 0.365 | 0.362 | 0.9 | 0.339 | 0.322 | 0.6 |
| 1836.0 | 9.667 | 10.071 | 0.5 | 9.836 | 9.503 | 0.5 | 9.192 | 9.696 | 0.1 | 3.739 | 3.290 | 0.1 |
| 1866.1 | 0.147 | 0.159 | 0.2 | 0.168 | 0.164 | 0.7 | 0.151 | 0.157 | 0.8 | 0.165 | 0.162 | 0.8 |
| 1907.1 | 0.418 | 0.450 | 0.4 | 0.466 | 0.473 | 0.9 | 0.392 | 0.384 | 0.8 | 0.633 | 0.630 | 1.0 |
| 1969.1 | 0.360 | 0.367 | 0.7 | 0.240 | 0.256 | 0.4 | 0.222 | 0.227 | 0.7 | 0.046 | 0.041 | 0.2 |
| 1988.1 | 3.776 | 3.626 | 0.4 | 3.055 | 3.162 | 0.5 | 3.435 | 3.472 | 0.9 | 4.556 | 4.142 | 0.1 |
| 1999.2 | 0.356 | 0.372 | 0.4 | 0.420 | 0.397 | 0.1 | 0.473 | 0.468 | 0.9 | 0.347 | 0.294 | **0.02*** |
| 2040.2 | 0.748 | 0.788 | 0.3 | 0.910 | 0.887 | 0.7 | 0.808 | 0.817 | 0.9 | 0.889 | 0.843 | 0.5 |
| 2070.2 | 0.409 | 0.410 | 1.0 | 0.474 | 0.476 | 1.0 | 0.526 | 0.523 | 0.9 | 1.315 | 1.275 | 0.7 |
| 2081.2 | 6.321 | 6.868 | 0.4 | 8.424 | 9.071 | 0.5 | 7.980 | 8.004 | 1.0 | 12.991 | 13.094 | 0.9 |
| 2111.2 | 0 | 0 | - | 0.077 | 0.081 | 0.6 | 0 | 0 | - | 0.084 | 0.072 | 0.2 |
| 2192.2 | 2.749 | 2.710 | 0.8 | 2.499 | 2.730 | 0.3 | 2.555 | 2.646 | 0.7 | 2.668 | 2.313 | 0.1 |
| 2203.2 | 0.247 | 0.253 | 0.6 | 0.262 | 0.255 | 0.6 | 0.271 | 0.268 | 0.8 | 0.435 | 0.407 | 0.3 |
| 2214.3 | 2.558 | 2.814 | 0.1 | 1.953 | 2.111 | 0.2 | 2.098 | 2.144 | 0.8 | 0.612 | 0.535 | 0.2 |
| 2244.3 | 0.764 | 0.802 | 0.6 | 1.033 | 1.070 | 0.7 | 1.128 | 1.069 | 0.6 | 1.967 | 1.738 | 0.3 |
| 2285.3 | 0.520 | 0.561 | 0.4 | 0.650 | 0.693 | 0.5 | 0.541 | 0.523 | 0.6 | 0.693 | 0.616 | 0.3 |
| 2326.3 | 0.104 | 0.112 | 0.4 | 0.089 | 0.100 | 0.2 | 0.111 | 0.100 | 0.3 | 0.340 | 0.327 | 0.7 |
| 2360.3 | 0.063 | 0.088 | 0.2 | 0.099 | 0.121 | 0.2 | 0.091 | 0.074 | 0.6 | 0.079 | 0.056 | **0.02*** |
| 2377.3 | 0.118 | 0.134 | 0.1 | 0.163 | 0.169 | 0.4 | 0.166 | 0.161 | 0.7 | 0.143 | 0.118 | **0.049*** |
| 2390.3 | 0.091 | 0.123 | 0.1 | 0.114 | 0.133 | 0.3 | 0.128 | 0.111 | 0.6 | 0.178 | 0.141 | 0.1 |
| 2396.3 | 2.311 | 2.244 | 0.8 | 2.080 | 2.444 | 0.1 | 2.343 | 2.316 | 0.9 | 3.206 | 2.591 | 0.1 |
| 2418.4 | 0.616 | 0.673 | 0.3 | 0.492 | 0.582 | 0.1 | 0.691 | 0.639 | 0.4 | 0.603 | 0.487 | 0.1 |
| 2431.4 | 0.067 | 0.094 | 0.2 | 0.067 | 0.092 | 0.1 | 0.088 | 0.082 | 0.9 | 0.087 | 0.068 | 0.1 |
| 2448.3 | 0.216 | 0.233 | 0.4 | 0.196 | 0.233 | 0.2 | 0.283 | 0.269 | 0.5 | 0.345 | 0.269 | 0.1 |
| 2459.4 | 3.988 | 4.467 | 0.2 | 3.873 | 4.598 | 0.1 | 4.338 | 4.050 | 0.4 | 5.386 | 5.152 | 0.7 |
| 2489.4 | 0.062 | 0.077 | **0.02*** | 0.083 | 0.093 | 0.2 | 0.073 | 0.071 | 0.8 | 0.084 | 0.061 | 0.1 |
| 2530.4 | 0.090 | 0.099 | 0.4 | 0.080 | 0.087 | 0.4 | 0.089 | 0.082 | 0.3 | 0.156 | 0.145 | 0.5 |
| 2547.4 | 0 | 0 | - | 0.049 | 0.050 | 0.8 | 0 | 0 | - | 0.069 | 0.050 | 0.3 |

| | | | | | | | | | | | | |
|---|---|---|---|---|---|---|---|---|---|---|---|---|
| 2564.4 | 0.048 | 0.071 | 0.1 | 0.080 | 0.097 | 0.2 | 0.080 | 0.068 | 0.7 | 0.086 | 0.059 | **0.01*** |
| 2592.5 | 0.298 | 0.368 | **0.015*** | 0.229 | 0.293 | **0.0098**** | 0.233 | 0.216 | 0.6 | 0.127 | 0.101 | 0.1 |
| 2600.5 | 0.039 | 0.040 | 0.9 | 0 | 0 | - | 0.116 | 0.097 | 0.7 | 0.049 | 0.042 | 0.4 |
| 2605.5 | 0.086 | 0.133 | 0.1 | 0.106 | 0.155 | 0.1 | 0 | 0 | - | 0.121 | 0.088 | **0.03*** |
| 2622.5 | 0.147 | 0.163 | 0.2 | 0.137 | 0.156 | 0.1 | 0.207 | 0.187 | 0.3 | 0.195 | 0.143 | 0.1 |
| 2635.5 | 0.078 | 0.104 | **0.035*** | 0.084 | 0.099 | 0.3 | 0.101 | 0.096 | 0.8 | 0.128 | 0.093 | **0.02*** |
| 2646.5 | 0.168 | 0.269 | 0.1 | 0.211 | 0.346 | 0.1 | 0.219 | 0.178 | 0.7 | 0.381 | 0.245 | **0.04*** |
| 2663.5 | 0.165 | 0.193 | 0.1 | 0.157 | 0.183 | 0.2 | 0.162 | 0.148 | 0.4 | 0.184 | 0.117 | 0.2 |
| 2704.5 | 0.305 | 0.366 | 0.2 | 0.202 | 0.267 | 0.1 | 0.305 | 0.271 | 0.2 | 0.593 | 0.538 | 0.6 |
| 2762.5 | 0 | 0 | - | 0.039 | 0.066 | 0.1 | 0 | 0 | - | 0 | 0 | - |
| 2779.6 | 0.110 | 0.187 | 0.1 | 0.114 | 0.179 | **0.04*** | 0.122 | 0.101 | 0.7 | 0.086 | 0.058 | **0.04*** |
| 2796.6 | 0.053 | 0.067 | **0.037*** | 0.057 | 0.074 | 0.1 | 0.073 | 0.065 | 0.6 | 0.058 | 0.036 | 0.1 |
| 2809.6 | 0.038 | 0.056 | **0.038*** | 0.048 | 0.063 | 0.1 | 0.057 | 0.051 | 0.8 | 0.057 | 0.034 | **0.004**** |
| 2837.6 | 1.176 | 1.496 | 0.1 | 0.857 | 1.172 | **0.046*** | 1.003 | 0.873 | 0.3 | 0.752 | 0.635 | 0.4 |
| 2867.6 | 0 | 0 | - | 0 | 0 | - | 0 | 0 | - | 0.037 | 0.021 | 0.1 |
| 2891.6 | 0.077 | 0.130 | 0.1 | 0.088 | 0.149 | 0.1 | 0.106 | 0.080 | 0.5 | 0.221 | 0.145 | 0.1 |
| 2908.6 | 0.057 | 0.070 | 0.1 | 0.053 | 0.066 | 0.2 | 0.068 | 0.058 | 0.3 | 0.108 | 0.064 | 0.3 |
| 2925.6 | 0.032 | 0.041 | 0.1 | 0.040 | 0.048 | 0.2 | 0 | 0 | - | 0.037 | 0.023 | 0.2 |
| 2966.6 | 0.054 | 0.092 | 0.1 | 0.075 | 0.131 | 0.1 | 0.074 | 0.072 | 1.0 | 0.080 | 0.047 | **0.03*** |
| 2983.6 | 0 | 0 | - | 0 | 0 | - | 0 | 0 | - | 0.030 | 0.014 | 0.2 |
| 2996.6 | 0 | 0 | - | 0 | 0 | - | 0 | 0 | - | 0.032 | 0.018 | **0.007**** |
| 3024.7 | 0.135 | 0.234 | 0.1 | 0.130 | 0.223 | 0.1 | 0.137 | 0.112 | 0.7 | 0.133 | 0.088 | **0.046*** |
| 3041.7 | 0.050 | 0.070 | **0.009**** | 0.059 | 0.074 | 0.1 | 0.061 | 0.056 | 0.7 | 0.055 | 0.028 | 0.2 |
| 3054.7 | 0 | 0 | - | 0 | 0 | - | 0 | 0 | - | 0.024 | 0.016 | **0.045*** |
| 3082.7 | 0.153 | 0.225 | **0.01*** | 0.130 | 0.182 | 0.1 | 0.193 | 0.161 | 0.2 | 0.271 | 0.218 | 0.3 |
| 3095.7 | 0 | 0 | - | 0 | 0 | - | 0 | 0 | - | 0.039 | 0.025 | **0.044*** |
| 3140.8 | 0.039 | 0.061 | **0.048*** | 0.042 | 0.070 | 0.1 | 0.050 | 0.047 | 0.8 | 0.032 | 0.024 | 0.2 |
| 3215.8 | 0.385 | 0.586 | **0.03*** | 0.538 | 0.732 | 0.1 | 0.394 | 0.361 | 0.6 | 0.124 | 0.079 | 0.2 |
| 3228.8 | 0 | 0 | - | 0.033 | 0.045 | 0.2 | 0 | 0 | - | 0.030 | 0.015 | 0.1 |
| 3269.8 | 0.104 | 0.222 | **0.02*** | 0.111 | 0.200 | 0.1 | 0.148 | 0.114 | 0.6 | 0.190 | 0.133 | 0.2 |
| 3286.9 | 0.029 | 0.044 | **0.01*** | 0.032 | 0.040 | 0.3 | 0.043 | 0.035 | 0.5 | 0.063 | 0.025 | 0.3 |
| 3327.9 | 0.031 | 0.053 | **0.049*** | 0.036 | 0.055 | 0.2 | 0.045 | 0.039 | 0.7 | 0.049 | 0.030 | **0.02*** |
| 3402.9 | 0.147 | 0.298 | **0.02*** | 0.228 | 0.374 | 0.1 | 0.185 | 0.160 | 0.7 | 0.081 | 0.048 | 0.1 |
| 3456.9 | 0.014 | 0.028 | **0.047*** | 0.020 | 0.033 | 0.2 | 0.023 | 0.020 | 0.8 | 0.037 | 0.022 | **0.03*** |
| 3460.9 | 0 | 0 | - | 0 | 0 | - | 0 | 0 | - | 0.098 | 0.064 | 0.1 |
| 3473.9 | 0 | 0 | - | 0 | 0 | - | 0 | 0 | - | 0.028 | 0.012 | 0.2 |
| 3514.8 | 0 | 0 | - | 0 | 0 | - | 0 | 0 | - | 0.018 | 0.012 | 0.2 |
| 3590.0 | 0.034 | 0.068 | 0.1 | 0.051 | 0.093 | 0.1 | 0.052 | 0.051 | 1.0 | 0.038 | 0.019 | 0.1 |
| 3631.0 | 0 | 0 | - | 0 | 0 | - | 0 | 0 | - | 0.019 | 0.012 | **0.04*** |
| 3648.0 | 0.069 | 0.155 | **0.01*** | 0.059 | 0.109 | 0.1 | 0.084 | 0.069 | 0.6 | 0.076 | 0.044 | 0.1 |
| 3764.1 | 0 | 0 | - | 0 | 0 | - | 0 | 0 | - | 0.023 | 0.011 | **0.01*** |
| 3777.1 | 0 | 0 | - | 0 | 0 | - | 0 | 0 | - | 0.021 | 0.011 | **0.03*** |
| 3835.2 | 0.021 | 0.044 | 0.1 | 0.020 | 0.038 | 0.1 | 0.026 | 0.024 | 0.9 | 0.029 | 0.016 | 0.1 |
| 3852.2 | 0 | 0 | - | 0 | 0 | - | 0 | 0 | - | 0.026 | 0.011 | 0.2 |
| 3893.2 | 0 | 0 | - | 0 | 0 | - | 0 | 0 | - | 0.025 | 0.014 | 0.1 |

| | | | | | | | | | | | | |
|---|---|---|---|---|---|---|---|---|---|---|---|---|
| 4009.2 | 0 | 0 | - | 0 | 0 | - | 0 | 0 | - | 0.020 | 0.009 | **0.009**** |
| 4026.3 | 0.060 | 0.142 | **0.02*** | 0.089 | 0.160 | 0.1 | 0.116 | 0.091 | 0.6 | 0.113 | 0.059 | 0.2 |
| 4039.3 | 0 | 0 | - | 0 | 0 | - | 0 | 0 | - | 0.018 | 0.007 | 0.1 |
| 4080.4 | 0 | 0 | - | 0 | 0 | - | 0 | 0 | - | 0.015 | 0.006 | **0.046*** |
| 4213.4 | 0.028 | 0.067 | **0.047*** | 0.034 | 0.066 | 0.1 | 0.059 | 0.051 | 0.7 | 0.063 | 0.030 | 0.1 |
| 4400.5 | 0 | 0 | - | 0 | 0 | - | 0.018 | 0.016 | 0.9 | 0.029 | 0.011 | **0.047*** |
| 4574.6 | 0 | 0 | - | 0 | 0 | - | 0 | 0 | - | 0.009 | 0.003 | **0.02*** |
| 4587.6 | 0 | 0 | - | 0 | 0 | - | 0 | 0 | - | 0.009 | 0.003 | **0.02*** |
| 4649.6 | 0 | 0 | - | 0 | 0 | - | 0 | 0 | - | 0.008 | 0.003 | 0.1 |

| *Female* | | | | | |
|---|---|---|---|---|---|
| **Cortex** | | | **Cerebellum** | | |
| CC | TT | *p-value* | CC | TT | *p-value* |
| 0.885 | 0.828 | 0.7 | 0.913 | 1.100 | 0.6 |
| 0.241 | 0.295 | 0.3 | 0.254 | 0.310 | 0.6 |
| 1.483 | 1.357 | 0.6 | 0.978 | 1.345 | 0.3 |
| 0.413 | 0.458 | 0.5 | 0.476 | 0.528 | 0.6 |
| 0.188 | 0.162 | 0.5 | 0.155 | 0.142 | 0.3 |
| 0.308 | 0.285 | 0.5 | 0.248 | 0.341 | 0.2 |
| 45.520 | 45.834 | 0.9 | 34.480 | 36.806 | 0.5 |
| 0.600 | 0.559 | 0.3 | 0.292 | 0.315 | 0.5 |
| 0.207 | 0.203 | 0.8 | 0.282 | 0.224 | 0.1 |
| 0.594 | 0.657 | 0.5 | 0.232 | 0.225 | 0.7 |
| 0.208 | 0.207 | 1.0 | 0.240 | 0.286 | 0.2 |
| 6.778 | 6.574 | 0.6 | 13.921 | 14.324 | 0.7 |
| 0.520 | 0.506 | 0.8 | 0.417 | 0.380 | 0.3 |
| 0.400 | 0.362 | 0.3 | 0.622 | 0.535 | 0.2 |
| 9.796 | 9.351 | 0.2 | 3.217 | 3.158 | 0.9 |
| 0.174 | 0.144 | 0.2 | 0.304 | 0.229 | 0.1 |
| 0.562 | 0.520 | 0.5 | 0.743 | 0.634 | 0.2 |
| 0.405 | 0.371 | **0.04*** | 0.076 | 0.071 | 0.4 |
| 3.161 | 3.118 | 0.8 | 4.772 | 4.679 | 0.6 |
| 0.425 | 0.427 | 1.0 | 0.502 | 0.469 | 0.6 |
| 0.909 | 0.869 | 0.6 | 1.147 | 1.017 | 0.4 |
| 0.430 | 0.445 | 0.8 | 1.952 | 1.578 | 0.2 |
| 6.616 | 6.879 | 0.7 | 9.477 | 8.896 | 0.6 |
| 0 | 0 | - | 0 | 0 | 0.7 |
| 2.238 | 2.337 | 0.6 | 3.138 | 2.945 | 0.5 |
| 0.268 | 0.249 | 0.5 | 0.661 | 0.622 | 0.6 |
| 2.476 | 2.299 | **0.04*** | 0.655 | 0.662 | 0.9 |
| 0.839 | 0.864 | 0.8 | 2.480 | 2.110 | 0.4 |
| 0.616 | 0.576 | 0.5 | 0.677 | 0.631 | 0.7 |
| 0.154 | 0.141 | 0.6 | 0.371 | 0.348 | 0.8 |
| 0.101 | 0.117 | 0.7 | 0.124 | 0.118 | 0.7 |
| 0.160 | 0.143 | 0.4 | 0.213 | 0.209 | 0.9 |
| 0.126 | 0.138 | 0.7 | 0.347 | 0.323 | 0.7 |
| 1.833 | 2.024 | 0.3 | 3.305 | 3.121 | 0.7 |
| 0.693 | 0.689 | 1.0 | 0.716 | 0.702 | 0.9 |
| 0.084 | 0.108 | 0.4 | 0.286 | 0.202 | 0.3 |
| 0.231 | 0.223 | 0.8 | 0.425 | 0.413 | 0.9 |
| 3.755 | 3.722 | 0.9 | 3.834 | 3.902 | 0.9 |
| 0.080 | 0.084 | 0.7 | 0.082 | 0.080 | 0.9 |
| 0.129 | 0.122 | 0.6 | 0.135 | 0.117 | 0.5 |
| 0 | 0 | - | 0 | 0 | 0.9 |

| | | | | | |
|---|---|---|---|---|---|
| 0.078 | 0.089 | 0.7 | 0.000 | 0.000 | - |
| 0.357 | 0.336 | 0.7 | 0.170 | 0.177 | 0.8 |
| 0 | 0 | - | 0 | 0 | 0.8 |
| 0.131 | 0.173 | 0.4 | 0.338 | 0.238 | 0.3 |
| 0.193 | 0.190 | 0.9 | 0.258 | 0.243 | 0.8 |
| 0.107 | 0.114 | 0.8 | 0.276 | 0.225 | 0.5 |
| 0.242 | 0.350 | 0.3 | 0.595 | 0.517 | 0.6 |
| 0.199 | 0.204 | 0.9 | 0.170 | 0.170 | 1.0 |
| 0.409 | 0.366 | 0.4 | 0.513 | 0.485 | 0.8 |
| 0 | 0 | - | 0 | 0 | - |
| 0.161 | 0.216 | 0.4 | 0.123 | 0.124 | 1.0 |
| 0.094 | 0.089 | 0.8 | 0.073 | 0.074 | 1.0 |
| 0.064 | 0.072 | 0.7 | 0.086 | 0.084 | 0.9 |
| 1.106 | 1.056 | 0.7 | 0.726 | 0.755 | 0.9 |
| 0 | 0 | - | 0 | 0 | - |
| 0.114 | 0.153 | 0.4 | 0.342 | 0.264 | 0.4 |
| 0.082 | 0.074 | 0.6 | 0.081 | 0.074 | 0.7 |
| 0 | 0 | - | 0 | 0 | - |
| 0.087 | 0.126 | 0.4 | 0.161 | 0.128 | 0.4 |
| 0 | 0 | - | 0 | 0 | 0.7 |
| 0 | 0 | - | 0 | 0 | 0.7 |
| 0.190 | 0.261 | 0.4 | 0.205 | 0.178 | 0.6 |
| 0.071 | 0.065 | 0.7 | 0.050 | 0.051 | 0.9 |
| 0 | 0 | - | 0 | 0 | 0.7 |
| 0.230 | 0.223 | 0.9 | 0.267 | 0.243 | 0.7 |
| 0 | 0 | - | 0 | 0 | 0.3 |
| 0.061 | 0.076 | 0.5 | 0.043 | 0.037 | 0.5 |
| 0.495 | 0.452 | 0.6 | 0.139 | 0.121 | 0.6 |
| 0 | 0 | - | 0 | 0 | 0.7 |
| 0.162 | 0.211 | 0.4 | 0.281 | 0.214 | 0.4 |
| 0.000 | 0.000 | - | 0.045 | 0.041 | 0.7 |
| 0.054 | 0.061 | 0.8 | 0.069 | 0.051 | 0.3 |
| 0.250 | 0.273 | 0.8 | 0.101 | 0.088 | 0.6 |
| 0.000 | 0.000 | - | 0.091 | 0.053 | 0.2 |
| 0 | 0 | 0.7 | 0 | 0 | 0.4 |
| 0 | 0 | - | 0 | 0 | 0.4 |
| 0 | 0 | - | 0 | 0 | 0.4 |
| 0.064 | 0.081 | 0.6 | 0.063 | 0.048 | 0.4 |
| 0 | 0 | - | 0 | 0 | 0.3 |
| 0.096 | 0.115 | 0.7 | 0.106 | 0.084 | 0.5 |
| 0 | 0 | - | 0 | 0 | 0.3 |
| 0 | 0 | - | 0 | 0 | 0.2 |
| 0.061 | 0.046 | 0.7 | 0.064 | 0.039 | 0.3 |
| 0 | 0 | - | 0 | 0 | 0.4 |
| 0 | 0 | - | 0 | 0 | 0.4 |

| | | | | | |
|---|---|---|---|---|---|
| 0 | 0 | - | 0 | 0 | 0.3 |
| 0.094 | 0.109 | 0.7 | 0.173 | 0.130 | 0.4 |
| 0 | 0 | 0.4 | 0 | 0 | 0.3 |
| 0 | 0 | - | 0 | 0 | 0.4 |
| 0.043 | 0.057 | 0.4 | 0.121 | 0.084 | 0.4 |
| 0 | 0 | - | 0 | 0 | 0.3 |
| 0 | 0 | - | 0 | 0 | 0.4 |
| 0 | 0 | - | 0 | 0 | 0.3 |
| 0 | 0 | - | 0 | 0 | 0.5 |

Table S2. Brain N-glycan structure, name, mass, and characteristics used in study.

| Glycan Name | *m/z* | 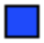 GlcNAc | 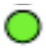 Mannose | 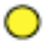 Galactose | 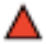 Fucose | 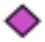 NeuAc | 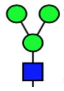 Pauci | 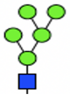 High-Man |
|---|---|---|---|---|---|---|---|---|
| F-Man-2 | 1141.6 | 2 | 2 | 0 | 1 | 0 | 1 | 0 |
| Man-3 | 1171.1 | 2 | 3 | 0 | 0 | 0 | 1 | 0 |
| F-Man-3 | 1345.8 | 2 | 3 | 0 | 1 | 0 | 1 | 0 |
| Man-4 | 1375.8 | 2 | 4 | 0 | 0 | 0 | 1 | 0 |
| A1 | 1416.8 | 3 | 3 | 0 | 0 | 0 | 0 | 0 |
| F-Man-4 | 1549.9 | 2 | 4 | 0 | 1 | 0 | 1 | 0 |
| Man-5 | 1579.9 | 2 | 5 | 0 | 0 | 0 | 0 | 1 |
| FA1 | 1590.9 | 3 | 3 | 0 | 1 | 0 | 0 | 0 |
| A1H4 | 1620.9 | 3 | 4 | 0 | 0 | 0 | 0 | 0 |
| A1B | 1661.9 | 4 | 3 | 0 | 0 | 0 | 0 | 0 |
| F-Man-5 | 1754.0 | 2 | 5 | 0 | 1 | 0 | 0 | 1 |
| Man-6 | 1784.0 | 2 | 6 | 0 | 0 | 0 | 0 | 1 |
| FA1H4 | 1795.0 | 3 | 4 | 0 | 1 | 0 | 0 | 0 |
| A1H5 | 1825.0 | 3 | 5 | 0 | 0 | 0 | 0 | 0 |
| FA1B | 1836.0 | 4 | 3 | 0 | 1 | 0 | 0 | 0 |
| A1BH4 | 1866.1 | 4 | 4 | 0 | 0 | 0 | 0 | 0 |
| A2B | 1907.1 | 5 | 3 | 0 | 0 | 0 | 0 | 0 |
| F2A1G1 | 1969.1 | 3 | 3 | 1 | 2 | 0 | 0 | 0 |
| Man-7 | 1988.1 | 2 | 7 | 0 | 0 | 0 | 0 | 1 |
| FA1H | 1999.2 | 3 | 5 | 0 | 1 | 0 | 0 | 0 |
| FA1BH4 | 2040.2 | 4 | 4 | 0 | 1 | 0 | 0 | 0 |
| A1BH5 | 2070.2 | 4 | 5 | 0 | 0 | 0 | 0 | 0 |
| FA2B | 2081.2 | 5 | 3 | 0 | 1 | 0 | 0 | 0 |
| A2BH4 | 2111.2 | 5 | 4 | 0 | 0 | 0 | 0 | 0 |
| Man-8 | 2192.2 | 2 | 8 | 0 | 0 | 0 | 0 | 1 |
| FA1G1H5 | 2203.2 | 3 | 5 | 1 | 1 | 0 | 0 | 0 |
| F2A1G1B | 2214.3 | 4 | 3 | 1 | 2 | 0 | 0 | 0 |
| FA1BH5 | 2244.3 | 4 | 5 | 0 | 1 | 0 | 0 | 0 |
| FA2BG1 | 2285.3 | 5 | 3 | 1 | 1 | 0 | 0 | 0 |
| FA3B | 2326.3 | 6 | 3 | 0 | 1 | 0 | 0 | 0 |
| FA1G1S1H4 | 2360.3 | 3 | 4 | 1 | 1 | 1 | 0 | 0 |
| F2A1G1H5 | 2377.3 | 3 | 5 | 1 | 2 | 0 | 0 | 0 |
| A1G1S1H5 | 2390.3 | 3 | 5 | 1 | 0 | 1 | 0 | 0 |
| Man-9 | 2396.3 | 2 | 9 | 0 | 0 | 0 | 0 | 1 |
| F2A1G1BH4 | 2418.4 | 4 | 4 | 1 | 2 | 0 | 0 | 0 |
| A1G1S1BH4 | 2431.4 | 4 | 4 | 1 | 0 | 1 | 0 | 0 |
| FAG1BH5 | 2448.3 | 4 | 5 | 1 | 1 | 0 | 0 | 0 |
| F2A1G1B | 2459.4 | 5 | 3 | 1 | 2 | 0 | 0 | 0 |
| FA2G1BH4 | 2489.4 | 5 | 4 | 1 | 1 | 0 | 0 | 0 |
| FA3G1B | 2530.4 | 6 | 3 | 1 | 1 | 0 | 0 | 0 |
| A1G1S2H4 | 2547.4 | 3 | 4 | 1 | 0 | 2 | 0 | 0 |
| FA1G1S1H5 | 2564.4 | 3 | 5 | 1 | 1 | 1 | 0 | 0 |
| F3A2G2 | 2592.5 | 4 | 3 | 2 | 3 | 0 | 0 | 0 |
| Man-9-G | 2600.5 | 2 | 9 | 0 | 0 | 0 | 0 | 1 |
| FA1G1S1BH4 | 2605.5 | 4 | 4 | 1 | 1 | 1 | 0 | 0 |
| F2A1G1BH5 | 2622.5 | 4 | 5 | 1 | 2 | 0 | 0 | 0 |
| A1G1S1BH5 | 2635.5 | 4 | 5 | 1 | 0 | 1 | 0 | 0 |
| FA2G1S1B | 2646.5 | 5 | 3 | 1 | 1 | 1 | 0 | 0 |
| F2A2G2B | 2663.5 | 5 | 3 | 2 | 2 | 0 | 0 | 0 |
| F2A3G1B | 2704.5 | 6 | 3 | 1 | 2 | 0 | 0 | 0 |
| FA1G1S2B | 2762.5 | 4 | 3 | 1 | 1 | 2 | 0 | 0 |
| F2A1G1S1BH4 | 2779.6 | 4 | 4 | 1 | 2 | 1 | 0 | 0 |
| F3A1G1BH5 | 2796.6 | 4 | 5 | 1 | 3 | 0 | 0 | 0 |
| FA1G1S1BH5 | 2809.6 | 4 | 5 | 1 | 1 | 1 | 0 | 0 |
| F3A2G2B | 2837.6 | 5 | 3 | 2 | 3 | 0 | 0 | 0 |
| F2A2G2BH4 | 2867.6 | 5 | 4 | 2 | 2 | 0 | 0 | 0 |

| | | | | | | | | |
|---|---|---|---|---|---|---|---|---|
| **FA3G1S1B** | **2891.6** | 6 | 3 | 1 | 1 | 1 | 0 | 0 |
| **F2A3G2B** | **2908.6** | 6 | 3 | 2 | 2 | 0 | 0 | 0 |
| **FA1G1S2H5** | **2925.6** | 3 | 5 | 1 | 1 | 2 | 0 | 0 |
| **FA2G2S2** | **2966.6** | 4 | 3 | 2 | 1 | 2 | 0 | 0 |
| **F2A1G1S1BH5** | **2983.6** | 4 | 5 | 1 | 2 | 1 | 0 | 0 |
| **A1G1S2BH** | **2996.6** | 4 | 5 | 1 | 0 | 2 | 0 | 0 |
| **F2A2G2S1B** | **3024.7** | 5 | 3 | 2 | 2 | 1 | 0 | 0 |
| **F3A3G3** | **3041.7** | 5 | 3 | 3 | 3 | 0 | 0 | 0 |
| **FA2G2S1BH4** | **3054.7** | 5 | 4 | 2 | 1 | 1 | 0 | 0 |
| **F3A3G2B** | **3082.7** | 6 | 3 | 2 | 3 | 0 | 0 | 0 |
| **FA3G2S1B** | **3095.7** | 6 | 3 | 2 | 1 | 1 | 0 | 0 |
| **F2A2G2S2** | **3140.8** | 4 | 3 | 2 | 2 | 2 | 0 | 0 |
| **F4A3G3** | **3215.8** | 5 | 3 | 3 | 4 | 0 | 0 | 0 |
| **F2A3G3S1** | **3228.8** | 5 | 3 | 3 | 2 | 1 | 0 | 0 |
| **F2A3G2S1B** | **3269.8** | 6 | 3 | 2 | 2 | 1 | 0 | 0 |
| **F3A3G3B** | **3286.9** | 6 | 3 | 3 | 3 | 0 | 0 | 0 |
| **FA2G2S3** | **3327.9** | 4 | 3 | 2 | 1 | 3 | 0 | 0 |
| **F3A2G2S1** | **3402.9** | 4 | 3 | 2 | 3 | 1 | 0 | 0 |
| **FA3G2S2B** | **3456.9** | 6 | 3 | 2 | 1 | 2 | 0 | 0 |
| **F4A3G3B** | **3460.9** | 6 | 3 | 3 | 4 | 0 | 0 | 0 |
| **F2A3G3S1B** | **3473.9** | 6 | 3 | 3 | 2 | 1 | 0 | 0 |
| **F2A4G2S1B** | **3514.8** | 7 | 3 | 2 | 2 | 1 | 0 | 0 |
| **F2A3G3S2** | **3590.0** | 5 | 3 | 3 | 2 | 2 | 0 | 0 |
| **F2A3G2S2B** | **3631.0** | 6 | 3 | 2 | 2 | 2 | 0 | 0 |
| **F3A3G3S1B** | **3648.0** | 6 | 3 | 3 | 3 | 1 | 0 | 0 |
| **F3A3G3S2** | **3764.1** | 5 | 3 | 3 | 3 | 2 | 0 | 0 |
| **FA3G3S3** | **3777.1** | 5 | 3 | 3 | 1 | 3 | 0 | 0 |
| **F2A3G3S2B** | **3835.2** | 6 | 3 | 3 | 2 | 2 | 0 | 0 |
| **F3A4G4S1** | **3852.2** | 6 | 3 | 4 | 3 | 1 | 0 | 0 |
| **F3A4G3S1B** | **3893.2** | 7 | 3 | 3 | 3 | 1 | 0 | 0 |
| **F3A3G3S2B** | **4009.2** | 6 | 3 | 3 | 3 | 2 | 0 | 0 |
| **F4A4G4S1** | **4026.3** | 6 | 3 | 4 | 4 | 1 | 0 | 0 |
| **F2A4G4S2** | **4039.3** | 6 | 3 | 4 | 2 | 2 | 0 | 0 |
| **F2A4G3S2B** | **4080.4** | 7 | 3 | 3 | 2 | 2 | 0 | 0 |
| **F3A4G4S2** | **4213.4** | 6 | 3 | 4 | 3 | 2 | 0 | 0 |
| **F2A4G4S3** | **4400.5** | 6 | 3 | 4 | 2 | 3 | 0 | 0 |
| **F3A4G4S3** | **4574.6** | 6 | 3 | 4 | 3 | 3 | 0 | 0 |
| **FA4G4S4** | **4587.6** | 6 | 3 | 4 | 1 | 4 | 0 | 0 |
| **F5A5G5S1** | **4649.6** | 7 | 3 | 5 | 5 | 1 | 0 | 0 |

| 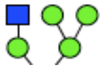 Hybrid | 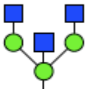 Bisecting | 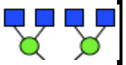 Antenna |
|---|---|---|
| 0 | 0 | 0 |
| 0 | 0 | 0 |
| 0 | 0 | 0 |
| 0 | 0 | 0 |
| 0 | 0 | 1 |
| 0 | 0 | 0 |
| 0 | 0 | 0 |
| 0 | 0 | 1 |
| 1 | 0 | 1 |
| 0 | 1 | 1 |
| 0 | 0 | 0 |
| 0 | 0 | 0 |
| 1 | 0 | 1 |
| 1 | 0 | 1 |
| 0 | 1 | 1 |
| 1 | 1 | 1 |
| 0 | 1 | 2 |
| 0 | 0 | 1 |
| 0 | 0 | 0 |
| 1 | 0 | 1 |
| 1 | 1 | 1 |
| 1 | 1 | 1 |
| 0 | 1 | 2 |
| 1 | 1 | 2 |
| 0 | 0 | 0 |
| 1 | 0 | 1 |
| 0 | 1 | 1 |
| 1 | 1 | 1 |
| 0 | 1 | 2 |
| 0 | 1 | 3 |
| 1 | 0 | 1 |
| 1 | 0 | 1 |
| 1 | 0 | 1 |
| 0 | 0 | 0 |
| 1 | 1 | 1 |
| 1 | 1 | 1 |
| 1 | 1 | 1 |
| 0 | 1 | 2 |
| 1 | 1 | 2 |
| 0 | 1 | 3 |
| 1 | 0 | 1 |
| 1 | 0 | 1 |
| 0 | 0 | 2 |
| 0 | 0 | 0 |
| 1 | 1 | 1 |
| 1 | 1 | 1 |
| 1 | 1 | 1 |
| 0 | 1 | 2 |
| 0 | 1 | 2 |
| 0 | 1 | 3 |
| 0 | 1 | 1 |
| 1 | 1 | 1 |
| 1 | 1 | 1 |
| 1 | 1 | 1 |
| 0 | 1 | 2 |
| 0 | 1 | 2 |

| | | |
|---|---|---|
| 0 | 1 | 3 |
| 0 | 1 | 3 |
| 1 | 0 | 1 |
| 0 | 0 | 2 |
| 1 | 1 | 1 |
| 1 | 1 | 1 |
| 0 | 1 | 2 |
| 0 | 0 | 3 |
| 1 | 1 | 2 |
| 0 | 1 | 3 |
| 0 | 1 | 3 |
| 0 | 0 | 2 |
| 0 | 0 | 3 |
| 0 | 0 | 3 |
| 0 | 1 | 3 |
| 0 | 1 | 3 |
| 0 | 0 | 2 |
| 0 | 0 | 2 |
| 0 | 1 | 3 |
| 0 | 1 | 3 |
| 0 | 1 | 3 |
| 0 | 1 | 4 |
| 0 | 0 | 3 |
| 0 | 1 | 3 |
| 0 | 1 | 3 |
| 0 | 0 | 3 |
| 0 | 0 | 3 |
| 0 | 1 | 3 |
| 0 | 0 | 4 |
| 0 | 1 | 4 |
| 0 | 1 | 3 |
| 0 | 0 | 4 |
| 0 | 0 | 4 |
| 0 | 1 | 4 |
| 0 | 0 | 4 |
| 0 | 0 | 4 |
| 0 | 0 | 4 |
| 0 | 0 | 4 |
| 0 | 0 | 5 |

**Table S3. Categorical analysis of protein N-glycans in 4-week old mice shows no difference in cortex and cerebellum*.**

| | Cortex | | | Cerebellum | | |
|---|---|---|---|---|---|---|
| **lycan Catego** | **CC** | **TT** | ***p-value*** | **CC** | **TT** | ***p-value*** |
| Paucimannose | 0.80 ± 0.03 | 0.77 ± 0.08 | 0.68 | 0.47 ± 0.04 | 0.46 ± 0.06 | 0.93 |
| High-mannose | 36.6 ± 1.25 | 36.2 ± 0.72 | 0.78 | 51.4 ± 1.53 | 49.8 ± 1.91 | 0.52 |
| lono-antennar | 20.4 ± 0.26 | 19.8 ± 0.57 | 0.36 | 13.8 ± 0.43 | 13.7 ± 0.46 | 0.93 |
| Bi-antennary | 26.7 ± 0.08 | 26.9 ± 0.35 | 0.48 | 27.3 ± 0.22 | 27.8 ± 0.38 | 0.30 |
| Tri-antennary | 12.9 ± 0.71 | 13.5 ± 0.54 | 0.53 | 5.97 ± 0.90 | 6.76 ± 1.02 | 0.58 |
| etra-antennar | 2.54 ± 0.35 | 2.75 ± 0.31 | 0.65 | 1.06 ± 0.32 | 1.50 ± 0.29 | 0.33 |
| Hybrid | 10.8 ± 0.37 | 10.4 ± 0.50 | 0.57 | 11.3 ± 0.40 | 11.2 ± 0.42 | 0.82 |
| Bisected | 50.6 ± 0.55 | 50.7 ± 0.44 | 0.85 | 44.0 ± 1.18 | 45.1 ± 1.47 | 0.58 |
| Galactose | 43.8 ± 1.36 | 44.6 ± 0.97 | 0.65 | 25.7 ± 1.81 | 27.3 ± 2.22 | 0.57 |
| Fucose | 60.6 ± 1.15 | 61.1 ± 0.75 | 0.72 | 44.8 ± 1.51 | 46.5 ± 1.89 | 0.51 |
| Sialic Acid | 14.8 ± 1.23 | 15.3 ± 0.80 | 0.73 | 6.17 ± 1.13 | 7.62 ± 1.06 | 0.37 |

*N = 5 per genotype and region, male mice only

Table S4. Brain O-glycan structure, name, mass, and characteristics used in study.

| Glycan Name | *m/z* | HexNAc | Hexose | NeuAc | NeuGc | Total Sialic Acid | Fucose |
|---|---|---|---|---|---|---|---|
| HexNAc Hex | 534.30 | 1 | 1 | 0 | 0 | 0 | 0 |
| HexNAc NeuAc | 691.41 | 1 | 0 | 1 | 0 | 1 | 0 |
| HexNAc2 Hex | 779.46 | 2 | 1 | 0 | 0 | 0 | 0 |
| HexNAc Hex NeuAc | 895.50 | 1 | 1 | 1 | 0 | 1 | 0 |
| HexNAc Hex2 Fuc | 912.51 | 1 | 2 | 0 | 0 | 0 | 1 |
| HexNAc Hex NeuGc | 925.53 | 1 | 1 | 0 | 1 | 1 | 0 |
| HexNAc2 Hex2 | 983.59 | 2 | 2 | 0 | 0 | 0 | 0 |
| HexNAc Hex NeuAc Fuc | 1069.63 | 1 | 1 | 1 | 0 | 1 | 1 |
| HexNAc Hex2 NeuAc | 1099.62 | 1 | 2 | 1 | 0 | 1 | 0 |
| HexNAc Hex NeuGc | 1129.65 | 1 | 2 | 0 | 1 | 1 | 0 |
| HexNAc2 Hex2 Fuc | 1157.66 | 2 | 2 | 0 | 0 | 0 | 1 |
| HexNAc Hex NeuAc2 | 1256.69 | 1 | 1 | 2 | 0 | 2 | 0 |
| HexNAc Hex NeuAc NeuGc | 1286.70 | 1 | 1 | 1 | 1 | 2 | 0 |
| HexNAc Hex NeuGc2 | 1316.72 | 1 | 1 | 0 | 2 | 2 | 0 |
| HexNAc2 Hex2 NeuAc | 1344.75 | 2 | 2 | 1 | 0 | 1 | 0 |
| HexNAc2 Hex3 Fuc | 1361.76 | 2 | 3 | 0 | 0 | 0 | 1 |
| HexNAc Hex2 NeuAc2 | 1460.80 | 1 | 2 | 2 | 0 | 2 | 0 |
| HexNAc2 Hex2 NeuAc Fuc | 1518.84 | 2 | 2 | 1 | 0 | 1 | 1 |
| HexNAc2 Hex3 Fuc2 | 1535.86 | 2 | 3 | 0 | 0 | 0 | 3 |
| HexNAc2 Hex3 NeuAc | 1548.85 | 2 | 3 | 1 | 0 | 1 | 0 |
| HexNAc2 Hex3 NeuGc | 1589.87 | 2 | 3 | 0 | 1 | 1 | 0 |
| HexNAc Hex NeuAc3 | 1617.88 | 1 | 1 | 3 | 0 | 3 | 0 |
| HexNAc2 Hex2 NeuAc2 | 1705.95 | 2 | 2 | 2 | 0 | 2 | 0 |
| HexNAc2 Hex3 NeuAc Fuc | 1722.97 | 2 | 3 | 1 | 0 | 1 | 1 |
| HexNAc2 Hex3 NeuAc2 | 1910.08 | 2 | 3 | 2 | 0 | 2 | 0 |
| HexNAc Hex NeuAc4 | 1979.12 | 1 | 1 | 4 | 0 | 4 | 0 |

| *Ser/Thr* O-Mannose | *Ser/Thr* O-GalNAc | Ambiguous |
|---|---|---|
| 0 | 0 | 1 |
| 0 | 1 | 0 |
| 0 | 1 | 0 |
| 0 | 1 | 0 |
| 1 | 0 | 0 |
| 0 | 1 | 0 |
| 0 | 1 | 0 |
| 0 | 1 | 0 |
| 1 | 0 | 0 |
| 1 | 0 | 0 |
| 0 | 1 | 0 |
| 0 | 1 | 0 |
| 0 | 1 | 0 |
| 0 | 1 | 0 |
| 0 | 0 | 1 |
| 1 | 0 | 0 |
| 1 | 0 | 0 |
| 0 | 0 | 1 |
| 1 | 0 | 0 |
| 1 | 0 | 0 |
| 1 | 0 | 0 |
| 0 | 1 | 0 |
| 0 | 1 | 0 |
| 1 | 0 | 0 |
| 1 | 0 | 0 |
| 0 | 1 | 0 |

**Table S5. Comparison of individual O-glycan abundance and categories in CC vs TT brain regions.**

| | *Male* | | | | | | |
|---|---|---|---|---|---|---|---|
| | **Cortex** | | | **Hippocampus** | | | |
| ***m/z*** | CC | TT | *p-value* | CC | TT | *p-value* | CC |
| 534.30 | 0.58 | 0.73 | 0.54 | 1.38 | 1.51 | 0.68 | 0.80 |
| 691.41 | 0.25 | 0.34 | 0.34 | 0.26 | 0.30 | 0.57 | 0.18 |
| 779.46 | 1.28 | 1.49 | 0.59 | 0.99 | 1.09 | 0.67 | 0.79 |
| 895.50 | 3.74 | 2.81 | 0.36 | 6.58 | 6.68 | 0.92 | 3.66 |
| 912.51 | 1.05 | 1.23 | 0.41 | 2.59 | 3.48 | **0.02*** | 1.40 |
| 925.53 | 0.13 | 0.16 | 0.33 | 0.00 | 0.00 | - | 0.15 |
| 983.59 | 0.39 | 0.48 | 0.46 | 0.28 | 0.46 | 0.06 | 0.29 |
| 1069.63 | 0.24 | 0.34 | 0.43 | 0.00 | 0.00 | - | 0.25 |
| 1099.62 | 5.70 | 6.14 | 0.55 | 11.43 | 9.38 | 0.06 | 7.21 |
| 1129.65 | 0.22 | 0.41 | 0.33 | 0.82 | 0.87 | 0.66 | 0.36 |
| 1157.66 | 5.95 | 6.55 | 0.64 | 2.90 | 4.58 | 0.07 | 3.18 |
| 1256.69 | 68.92 | 66.46 | 0.08 | 64.54 | 62.85 | 0.15 | 68.65 |
| 1286.70 | 0.69 | 0.50 | 0.28 | 0.43 | 0.55 | 0.28 | 1.13 |
| 1316.72 | 0.18 | 0.25 | 0.29 | 0.00 | 0.00 | - | 0.23 |
| 1344.75 | 3.28 | 2.90 | 0.38 | 1.70 | 1.62 | 0.68 | 2.41 |
| 1361.76 | 0.44 | 0.60 | 0.09 | 0.41 | 0.48 | 0.47 | 0.67 |
| 1460.80 | 1.29 | 1.62 | 0.30 | 1.41 | 1.34 | 0.59 | 1.40 |
| 1518.84 | 1.33 | 1.32 | 0.96 | 0.31 | 0.45 | 0.14 | 0.59 |
| 1535.86 | 0.53 | 0.70 | 0.17 | 0.52 | 0.60 | 0.47 | 0.52 |
| 1548.85 | 0.55 | 0.65 | 0.47 | 0.61 | 0.61 | 0.96 | 0.79 |
| 1589.87 | 0.08 | 0.14 | **0.006**** | 0.13 | 0.14 | 0.85 | 0.23 |
| 1617.88 | 1.50 | 1.88 | 0.24 | 1.29 | 1.46 | 0.46 | 3.05 |
| 1705.95 | 0.77 | 0.85 | 0.44 | 0.28 | 0.35 | 0.17 | 0.60 |
| 1722.97 | 0.33 | 0.48 | **0.03*** | 0.33 | 0.40 | 0.39 | 0.42 |
| 1910.08 | 0.49 | 0.81 | 0.06 | 0.73 | 0.72 | 0.91 | 0.85 |
| 1979.12 | 0.10 | 0.19 | 0.06 | 0.07 | 0.09 | 0.41 | 0.18 |

| | | *Male* | | | | | | |
|---|---|---|---|---|---|---|---|---|
| | | **Cortex** | | | **Hippocampus** | | | |
| **Glycan Category** | | CC | TT | *p-value* | CC | TT | *p-value* | CC |
| *All O-glycans* | O-GalNAc | 84.1 ± 0.3 | 82.3 ± 0.4 | **0.01*** | 77.6 ± 0.2 | 78.4 ± 0.9 | 0.4 | 82 ± 1 |
| | O-Mannose | 10.7 ± 0.4 | 12.8 ± 0.6 | **0.03*** | 19.0 ± 0.3 | 18.0 ± 0.6 | 0.2 | 14 ± 1 |
| | Fucose | 9.9 ± 1.1 | 11.2 ± 1.3 | 0.4 | 7.1 ± 0.3 | 10.0 ± 1.1 | 0.07 | 7.0 ± 1.1 |
| | NeuAc | 89.2 ± 0.9 | 87.3 ± 1.1 | 0.2 | 90.0 ± 0.2 | 86.8 ± 1.3 | 0.09 | 91.4 ± 1.2 |
| | NeuGc | 1.3 ± 0.2 | 1.5 ± 0.3 | 0.6 | 1.39 ± 0.01 | 1.56 ± 0.2 | 0.4 | 2.10 ± 0.08 |
| *O-GalNAc* | Core 1 | 90 ± 1 | 89 ± 1 | 0.4 | 94.3 ± 0.1 | 91.7 ± 1.1 | 0.09 | 94.1 ± 1.1 |
| | Core 2 | 10 ± 1 | 11 ± 1 | 0.4 | 5.7 ± 0.1 | 8.3 ± 1.1 | 0.09 | 5.9 ± 1.1 |
| | Fucose | 7.4 ± 1.0 | 8.4 ± 1.2 | 0.5 | 3.7 ± 0.2 | 5.9 ± 0.9 | 0.08 | 4.2 ± 1.1 |
| | NeuAc | 91 ± 1 | 89 ± 1 | 0.3 | 94.6 ± 0.1 | 92.2 ± 1.0 | 0.09 | 94.3 ± 1.2 |
| | NeuGc | 1.2 ± 0.2 | 1.1 ± 2 | 0.8 | 0.56 ± 0.06 | 0.70 ± 0.1 | 0.3 | 1.8 ± 0.2 |
| *O-Mannose* | Core M1 | 77 ± 2 | 74 ± 2 | 0.3 | 86 ± 1 | 84 ± 2 | 0.5 | 74 ± 4 |
| | Core M2 | 23 ± 2 | 26 ± 2 | 0.3 | 15 ± 1 | 16 ± 2 | 0.5 | 26 ± 4 |
| | Fucose | 22 ± 1 | 24 ± 2 | 0.5 | 20.3 ± 0.7 | 27.4 ± 1.6 | **0.02*** | 22 ± 2 |
| | NeuAc | 78 ± 1 | 76 ± 2 | 0.3 | 76.5 ± 0.5 | 69.2 ± 1.7 | **0.02*** | 77 ± 2 |
| | NeuGc | 2.8 ± 0.6 | 4.2 ± 1.3 | 0.3 | 0.05 ± 0.001 | 0.06 ± 0.006 | 0.4 | 4.3 ± 0.5 |

| | | | | | | *Female* | |
|---|---|---|---|---|---|---|---|
| **Striatum** | | **Cerebellum** | | | **Cortex** | | |
| TT | *p-value* | CC | TT | *p-value* | CC | TT | *p-value* |
| 0.99 | 0.34 | 0.54 | 0.98 | 0.33 | 1.26 | 2.56 | 0.27 |
| 0.18 | 0.98 | 0.53 | 0.76 | 0.26 | 1.22 | 1.30 | 0.92 |
| 0.98 | 0.65 | 2.44 | 4.24 | 0.24 | 3.48 | 5.60 | 0.43 |
| 4.10 | 0.52 | 7.20 | 6.77 | 0.81 | 5.74 | 5.52 | 0.80 |
| 1.31 | 0.80 | 4.64 | 3.66 | 0.41 | 1.51 | 1.81 | 0.68 |
| 0.13 | 0.47 | 0.36 | 0.36 | 0.98 | 0.19 | 0.24 | 0.49 |
| 0.22 | 0.44 | 0.39 | 0.45 | 0.68 | 0.36 | 0.51 | 0.26 |
| 0.24 | 0.89 | 0.79 | 1.29 | 0.37 | 1.26 | 1.27 | 0.99 |
| 6.90 | 0.81 | 18.38 | 20.86 | 0.34 | 7.94 | 7.82 | 0.88 |
| 0.32 | 0.69 | 0.47 | 0.48 | 0.93 | 0.34 | 0.91 | 0.29 |
| 2.73 | 0.64 | 4.03 | 2.91 | 0.29 | 4.35 | 5.76 | 0.32 |
| 69.06 | 0.85 | 55.20 | 52.79 | 0.54 | 63.22 | 56.14 | 0.29 |
| 1.02 | 0.62 | 1.07 | 0.74 | 0.06 | 0.42 | 0.42 | 0.95 |
| 0.20 | 0.53 | 0.14 | 0.12 | 0.57 | 0.20 | 0.39 | 0.10 |
| 2.67 | 0.43 | 0.74 | 0.71 | 0.61 | 3.01 | 3.51 | 0.44 |
| 0.49 | 0.17 | 0.13 | 0.12 | 0.82 | 0.36 | 0.78 | 0.13 |
| 1.44 | 0.86 | 0.73 | 0.74 | 0.95 | 1.15 | 0.92 | 0.51 |
| 0.63 | 0.75 | 0.26 | 0.21 | 0.31 | 0.72 | 0.88 | 0.58 |
| 0.60 | 0.55 | 0.10 | 0.10 | 0.95 | 0.36 | 0.60 | 0.15 |
| 0.78 | 0.93 | 0.16 | 0.15 | 0.82 | 0.44 | 0.55 | 0.44 |
| 0.21 | 0.58 | 0.11 | 0.10 | 0.91 | 0.11 | 0.13 | 0.55 |
| 2.77 | 0.72 | 0.94 | 0.84 | 0.33 | 1.15 | 1.03 | 0.64 |
| 0.58 | 0.86 | 0.32 | 0.27 | 0.18 | 0.53 | 0.52 | 0.93 |
| 0.43 | 0.95 | 0.09 | 0.08 | 0.91 | 0.23 | 0.27 | 0.60 |
| 0.87 | 0.92 | 0.16 | 0.16 | 0.88 | 0.38 | 0.39 | 0.98 |
| 0.16 | 0.66 | 0.07 | 0.09 | 0.58 | 0.06 | 0.18 | 0.26 |

| | | | | | | *Female* | |
|---|---|---|---|---|---|---|---|
| **Striatum** | | **Cerebellum** | | | **Cortex** | | |
| TT | *p-value* | CC | TT | *p-value* | CC | TT | *p-value* |
| 82 ± 1 | 1.0 | 74 ± 2 | 72 ± 3 | 0.6 | 82 ± 0.3 | 79 ± 2 | 0.23 |
| 13 ± 1 | 0.7 | 25 ± 3 | 27 ± 3 | 0.7 | 13 ± 0.004 | 14 ± 2 | 0.47 |
| 6.4 ± 0.5 | 0.7 | 10 ± 2 | 8 ± 1 | 0.5 | 8.8 ± 0.1 | 11 ± 2 | 0.21 |
| 91.8 ± 0.6 | 0.8 | 87 ± 2 | 86 ± 2 | 1.0 | 87.5 ± 0.8 | 81 ± 5 | 0.28 |
| 1.87 ± 0.07 | 0.1 | 2.2 ± 0.2 | 1.82 ± 0.07 | 0.1 | 1.27 ± 0.08 | 2.1 ± 0.5 | 0.21 |
| 94.5 ± 0.3 | 0.7 | 90 ± 1 | 89 ± 2 | 0.6 | 89.4 ± 0.2 | 84 ± 4 | 0.26 |
| 5.5 ± 0.3 | 0.7 | 10 ± 1 | 11 ± 2 | 0.6 | 10.6 ± 0.2 | 16 ± 4 | 0.26 |
| 3.6 ± 0.3 | 0.6 | 7 ± 1 | 6 ± 1 | 0.7 | 6.8 ± 0.1 | 9 ± 1 | 0.20 |
| 94.8 ± 0.3 | 0.7 | 90 ± 1 | 89 ± 2 | 0.6 | 89.6 ± 0.3 | 84 ± 4 | 0.26 |
| 1.6 ± 0.1 | 0.4 | 2.1 ± 0.1 | 1.7 ± 0.1 | **0.03*** | 0.99 ± 0.05 | 1.3 ± 0.1 | 0.07 |
| 75 ± 3 | 0.8 | 97.0 ± 0.6 | 97.4 ± 0.3 | 0.6 | 85 ± 1 | 80 ± 3 | 0.16 |
| 25 ± 3 | 0.8 | 3.0 ± 0.6 | 2.6 ±0.3 | 0.6 | 15 ± 1 | 20 ± 3 | 0.16 |
| 21 ± 2 | 0.8 | 19 ± 2 | 15 ± 2 | 0.1 | 19.2 ± 0.8 | 23 ± 4 | 0.35 |
| 78 ± 2 | 0.6 | 79 ± 2 | 84 ± 2 | 0.2 | 79 ± 2 | 72 ± 5 | 0.30 |
| 3.9 ± 0.3 | 0.5 | 2.3 ± 0.3 | 2.2 ± 0.2 | 0.7 | 3.5 ± 0.9 | 7 ± 2 | 0.23 |

**Table S6. Differentially expressed genes in cortex and cerebellum of CC vs TT mice at False Discovery Rate (FDR) of 0.05**

| Cortex | Cerebellum | | | |
|---|---|---|---|---|
| Btg2 | Accn4 | Egr3 | Kcnk15 | Slc16a8 |
| Cyr61 | Alox5 | Fanci | Ldb2 | Sst |
| Npas4 | Ankrd34c | Gda | Lhx2 | Stk32b |
| | Bcl11b | Gm1337 | Lhx9 | Tcf7l2 |
| | Camkv | Gpr26 | Map3k15 | Tfap2d |
| | Cbln2 | Grp | Meis2 | Tmem215 |
| | Chrm3 | Hs3st4 | Npas4 | Tox3 |
| | Cntn5 | Kcnc2 | Nrgn | Vip |
| | Col8a1 | Kcnh5 | Pde11a | Zfhx3 |
| | Cpne5 | Kcnip2 | Ptgs2 | Zfp831 |
| | Dlgap2 | Kcnj2 | Slc15a2 | |

**Table S7. N-glycoprotein levels in cortex of TT mice relative to CC controls.**

| Symbol | Protein Accessions | PSMs | Glycosite | FC | logFC | P.Value | logPval | adj.P.Val |
|---|---|---|---|---|---|---|---|---|
| Slc1a2 | P43006 | 6 | 6 | -4.897 | -2.307 | 0.001 | 3.246 | 0.004 |
| Gde1 | Q9JL56 | 2 | 2 | -3.878 | -1.979 | 0.000 | 3.758 | 0.002 |
| Slc7a1 | Q09143 | 1 | 1 | -3.836 | -1.963 | 0.001 | 3.251 | 0.004 |
| Astn2 | Q80Z10 | 1 | 1 | -3.575 | -1.865 | 0.000 | 4.239 | 0.001 |
| Panx1 | Q9JIP4 | 1 | 1 | -3.367 | -1.782 | 0.003 | 2.560 | 0.013 |
| Hsd11b1 | P50172 | 1 | 1 | -3.241 | -1.730 | 0.001 | 2.985 | 0.006 |
| Epdr1 | Q99M71 | 2 | 2 | -2.955 | -1.603 | 0.012 | 1.916 | 0.035 |
| Glg1 | Q61543 | 2 | 2 | -2.946 | -1.599 | 0.000 | 3.574 | 0.003 |
| Serpina10 | Q8R121 | 1 | 1 | -2.762 | -1.511 | 0.010 | 1.991 | 0.031 |
| Nptn | P97300 | 8 | 8 | -2.658 | -1.459 | 0.000 | 4.299 | 0.001 |
| Tenm3 | Q9WTS6 | 2 | 2 | -2.653 | -1.456 | 0.003 | 2.506 | 0.014 |
| Tf | Q921I1 | 1 | 1 | -2.609 | -1.434 | 0.000 | 3.704 | 0.002 |
| Chl1 | P70232 | 6 | 6 | -2.512 | -1.383 | 0.000 | 3.390 | 0.004 |
| Ncam1 | P13595 | 4 | 4 | -2.505 | -1.379 | 0.001 | 2.900 | 0.007 |
| Tmeff1 | Q6PFE7 | 2 | 2 | -2.464 | -1.357 | 0.000 | 3.530 | 0.003 |
| Itgb1 | P09055 | 3 | 3 | -2.461 | -1.355 | 0.000 | 5.642 | 0.000 |
| Reln | Q60841 | 1 | 1 | -2.421 | -1.334 | 0.001 | 3.005 | 0.006 |
| Enpep | P16406 | 3 | 3 | -2.367 | -1.304 | 0.001 | 3.000 | 0.006 |
| Atp1a1 | Q8VDN2 | 1 | 1 | -2.300 | -1.266 | 0.008 | 2.093 | 0.027 |
| Atp1b1 | P14094 | 7 | 7 | -2.295 | -1.262 | 0.000 | 3.594 | 0.003 |
| Ano6 | Q6P9J9 | 1 | 1 | -2.228 | -1.224 | 0.000 | 3.957 | 0.002 |
| Slc44a1 | Q6X893 | 2 | 2 | -2.214 | -1.215 | 0.005 | 2.334 | 0.018 |
| Nrp1 | P97333 | 2 | 2 | -2.204 | -1.209 | 0.001 | 3.289 | 0.004 |
| Prnp | P04925 | 3 | 3 | -2.164 | -1.185 | 0.000 | 3.925 | 0.002 |
| B4galnt1 | Q09200 | 1 | 1 | -2.105 | -1.149 | 0.010 | 2.017 | 0.031 |
| Gabbr1 | Q9WV18 | 3 | 3 | -1.974 | -1.066 | 0.004 | 2.396 | 0.017 |
| Cd47 | Q61735 | 3 | 4 | -1.941 | -1.044 | 0.001 | 3.235 | 0.004 |
| Nfasc | Q810U3 | 7 | 7 | -1.920 | -1.030 | 0.004 | 2.369 | 0.017 |
| Cdh5 | P55284 | 1 | 1 | -1.885 | -1.007 | 0.001 | 3.269 | 0.004 |
| Sema4d | O09126 | 3 | 3 | -1.885 | -1.007 | 0.003 | 2.542 | 0.013 |
| Atp1b3 | P97370 | 2 | 2 | -1.747 | -0.911 | 0.005 | 2.318 | 0.018 |
| Slc12a6 | Q924N4 | 2 | 2 | -1.711 | -0.885 | 0.001 | 3.016 | 0.006 |
| Lamc1 | P02468 | 3 | 3 | -1.709 | -0.883 | 0.010 | 1.989 | 0.031 |
| Stt3b | Q3TDQ1 | 2 | 2 | -1.674 | -0.858 | 0.017 | 1.761 | 0.045 |
| Hyou1 | Q9JKR6 | 4 | 4 | -1.668 | -0.853 | 0.015 | 1.817 | 0.041 |
| Gpr158 | Q8C419 | 1 | 1 | -1.619 | -0.817 | 0.020 | 1.704 | 0.050 |
| Pcdh19 | Q80TF3 | 1 | 1 | -1.578 | -0.785 | 0.017 | 1.775 | 0.044 |
| Cst3 | P21460 | 1 | 1 | -1.553 | -0.765 | 0.086 | 1.067 | 0.156 |
| Sorl1 | O88307 | 5 | 5 | -1.553 | -0.765 | 0.001 | 3.298 | 0.004 |
| Slco1c1 | Q9ERB5 | 1 | 1 | -1.547 | -0.761 | 0.000 | 3.457 | 0.003 |
| Plxdc1 | Q91ZV7 | 1 | 1 | -1.546 | -0.760 | 0.021 | 1.675 | 0.052 |
| Scg3 | P47867 | 1 | 1 | -1.512 | -0.733 | 0.012 | 1.923 | 0.035 |

| | | | | | | | | |
|---|---|---|---|---|---|---|---|---|
| Tmem30b | Q8BHG3 | 1 | 1 | -1.509 | -0.731 | 0.001 | 2.835 | 0.007 |
| Ceacam1 | P31809 | 2 | 2 | -1.479 | -0.707 | 0.041 | 1.387 | 0.089 |
| ef1akmt4-Ece | P0DPD9 | 1 | 1 | -1.414 | -0.654 | 0.054 | 1.268 | 0.111 |
| Smpd1 | Q04519 | 1 | 1 | -1.395 | -0.638 | 0.021 | 1.674 | 0.052 |
| Ero1b | Q8R2E9 | 2 | 3 | -1.395 | -0.637 | 0.010 | 2.003 | 0.031 |
| Itih3 | Q61704 | 1 | 1 | -1.392 | -0.635 | 0.109 | 0.962 | 0.190 |
| Enpp5 | Q9EQG7 | 1 | 1 | -1.366 | -0.613 | 0.245 | 0.612 | 0.345 |
| Slc38a3 | Q9DCP2 | 1 | 2 | -1.349 | -0.599 | 0.031 | 1.513 | 0.072 |
| Grin2b | Q01097 | 6 | 6 | -1.337 | -0.588 | 0.003 | 2.574 | 0.013 |
| Slc3a2 | P10852 | 3 | 3 | -1.316 | -0.570 | 0.001 | 2.879 | 0.007 |
| Slc12a5 | Q91V14 | 3 | 3 | -1.311 | -0.566 | 0.058 | 1.233 | 0.118 |
| Emb | P21995 | 7 | 9 | -1.302 | -0.557 | 0.008 | 2.111 | 0.026 |
| Plxnb2 | B2RXS4 | 3 | 3 | -1.287 | -0.544 | 0.068 | 1.170 | 0.131 |
| Adgre5 | Q9Z0M6 | 1 | 1 | -1.283 | -0.540 | 0.069 | 1.163 | 0.132 |
| Sel1l | Q9Z2G6 | 1 | 1 | -1.263 | -0.523 | 0.102 | 0.990 | 0.179 |
| Ephb2 | P54763 | 2 | 2 | -1.244 | -0.506 | 0.014 | 1.844 | 0.040 |
| Ntrk2 | P15209 | 8 | 8 | -1.242 | -0.504 | 0.014 | 1.848 | 0.040 |
| Spock2 | Q9ER58 | 2 | 2 | -1.237 | -0.500 | 0.004 | 2.452 | 0.015 |
| Apoh | Q01339 | 2 | 2 | -1.221 | -0.485 | 0.171 | 0.766 | 0.265 |
| Nt5e | Q61503 | 2 | 2 | -1.217 | -0.482 | 0.022 | 1.667 | 0.053 |
| Serpini1 | O35684 | 2 | 2 | -1.213 | -0.477 | 0.035 | 1.458 | 0.079 |
| Itgb2 | P11835 | 1 | 1 | -1.212 | -0.476 | 0.067 | 1.173 | 0.131 |
| Atrn | Q9WU60 | 3 | 4 | -1.189 | -0.455 | 0.012 | 1.918 | 0.035 |
| Adgrb3 | Q80ZF8 | 4 | 4 | -1.151 | -0.420 | 0.026 | 1.587 | 0.063 |
| Pzp | Q61838 | 4 | 4 | -1.141 | -0.411 | 0.118 | 0.928 | 0.201 |
| Ace | P09470 | 2 | 2 | -1.138 | -0.407 | 0.036 | 1.440 | 0.080 |
| Ptgfrn | Q9WV91 | 2 | 2 | -1.122 | -0.393 | 0.085 | 1.071 | 0.156 |
| Il1rapl1 | P59823 | 1 | 1 | -1.122 | -0.392 | 0.209 | 0.680 | 0.313 |
| Cntn3 | Q07409 | 5 | 5 | -1.103 | -0.374 | 0.128 | 0.894 | 0.216 |
| Cxadr | P97792 | 1 | 1 | -1.103 | -0.374 | 0.242 | 0.617 | 0.343 |
| Lamp1 | P11438 | 5 | 7 | -1.097 | -0.368 | 0.072 | 1.143 | 0.136 |
| Slc39a6 | Q8C145 | 2 | 2 | -1.058 | -0.330 | 0.118 | 0.929 | 0.201 |
| Ano3 | A2AHL1 | 2 | 2 | -1.057 | -0.329 | 0.210 | 0.678 | 0.313 |
| Lama5 | Q61001 | 3 | 3 | -1.050 | -0.322 | 0.438 | 0.359 | 0.547 |
| Psap | Q61207 | 4 | 4 | -1.044 | -0.316 | 0.060 | 1.219 | 0.121 |
| Mdga2 | P60755 | 1 | 1 | -1.041 | -0.312 | 0.324 | 0.489 | 0.431 |
| Scn1a | A2APX8 | 1 | 1 | -1.027 | -0.299 | 0.053 | 1.280 | 0.110 |
| Ap2b1 | Q9DBG3 | 1 | 1 | -1.017 | -0.288 | 0.361 | 0.443 | 0.471 |
| Slitrk1 | Q810C1 | 2 | 2 | -1.015 | -0.287 | 0.460 | 0.337 | 0.571 |
| Sort1 | Q6PHU5 | 2 | 2 | -1.013 | -0.285 | 0.217 | 0.664 | 0.316 |
| Grin1 | P35438 | 6 | 6 | -1.013 | -0.285 | 0.168 | 0.776 | 0.263 |
| Thy1 | P01831 | 2 | 2 | -1.011 | -0.283 | 0.392 | 0.407 | 0.505 |
| Gria2 | P23819 | 2 | 2 | -1.011 | -0.282 | 0.141 | 0.850 | 0.233 |
| Elapor2 | Q3UZV7 | 1 | 2 | -0.998 | -0.270 | 0.364 | 0.439 | 0.472 |
| Lrp1 | Q91ZX7 | 11 | 11 | -0.996 | -0.268 | 0.265 | 0.577 | 0.368 |

| | | | | | | | | |
|---|---|---|---|---|---|---|---|---|
| Clu | Q06890 | 4 | 4 | -0.990 | -0.261 | 0.255 | 0.593 | 0.357 |
| Ppt1 | O88531 | 3 | 3 | -0.954 | -0.224 | 0.169 | 0.773 | 0.263 |
| Enpp4 | Q8BTJ4 | 2 | 2 | -0.954 | -0.224 | 0.214 | 0.670 | 0.314 |
| Alcam | Q61490 | 8 | 8 | -0.949 | -0.219 | 0.207 | 0.684 | 0.313 |
| Tnr | Q8BYI9 | 8 | 9 | -0.944 | -0.214 | 0.139 | 0.857 | 0.232 |
| Ptprj | Q64455 | 3 | 5 | -0.938 | -0.208 | 0.403 | 0.395 | 0.513 |
| Hp | Q61646 | 2 | 2 | -0.936 | -0.205 | 0.600 | 0.222 | 0.696 |
| Atp1b2 | P14231 | 7 | 8 | -0.935 | -0.204 | 0.306 | 0.515 | 0.411 |
| Slc2a3 | P32037 | 1 | 1 | -0.933 | -0.202 | 0.329 | 0.483 | 0.432 |
| Serpina6 | Q06770 | 2 | 2 | -0.932 | -0.201 | 0.238 | 0.624 | 0.340 |
| Slc6a6 | O35316 | 1 | 1 | -0.905 | -0.172 | 0.492 | 0.308 | 0.604 |
| B3glct | Q8BHT6 | 1 | 1 | -0.900 | -0.166 | 0.588 | 0.231 | 0.690 |
| Omg | Q63912 | 2 | 2 | -0.894 | -0.160 | 0.594 | 0.226 | 0.693 |
| Serpinc1 | P32261 | 2 | 2 | -0.877 | -0.142 | 0.532 | 0.274 | 0.642 |
| Serpina3m | Q03734 | 1 | 1 | -0.867 | -0.132 | 0.580 | 0.237 | 0.688 |
| Mag | P20917 | 6 | 8 | -0.833 | -0.094 | 0.686 | 0.164 | 0.792 |
| Entpd2 | O55026 | 2 | 2 | -0.829 | -0.089 | 0.747 | 0.126 | 0.848 |
| Lama2 | Q60675 | 7 | 7 | -0.827 | -0.088 | 0.566 | 0.247 | 0.677 |
| Tmem30a | Q8VEK0 | 1 | 2 | -0.827 | -0.087 | 0.568 | 0.246 | 0.677 |
| Adcy3 | Q8VHH7 | 1 | 1 | -0.825 | -0.085 | 0.776 | 0.110 | 0.857 |
| Cdh13 | Q9WTR5 | 3 | 3 | -0.811 | -0.069 | 0.845 | 0.073 | 0.891 |
| Cntn2 | Q61330 | 3 | 4 | -0.809 | -0.067 | 0.769 | 0.114 | 0.857 |
| Tmem87a | Q8BXN9 | 3 | 5 | -0.807 | -0.065 | 0.827 | 0.082 | 0.891 |
| Cdh10 | P70408 | 1 | 1 | -0.803 | -0.060 | 0.734 | 0.134 | 0.838 |
| Lgi1 | Q9JIA1 | 2 | 2 | -0.794 | -0.051 | 0.778 | 0.109 | 0.857 |
| Lamp5 | Q9D387 | 2 | 2 | -0.794 | -0.050 | 0.780 | 0.108 | 0.857 |
| Cd82 | P40237 | 4 | 5 | -0.792 | -0.048 | 0.795 | 0.100 | 0.865 |
| Lgals3bp | Q07797 | 2 | 2 | -0.782 | -0.036 | 0.869 | 0.061 | 0.913 |
| Dpp10 | Q6NXK7 | 4 | 5 | -0.775 | -0.029 | 0.926 | 0.033 | 0.964 |
| Lgi2 | Q8K4Z0 | 2 | 2 | -0.763 | -0.014 | 0.927 | 0.033 | 0.964 |
| Ptprz1 | B9EKR1 | 10 | 10 | -0.760 | -0.012 | 0.937 | 0.028 | 0.969 |
| Cdh2 | P15116 | 3 | 3 | -0.755 | -0.005 | 0.986 | 0.006 | 0.992 |
| Gfra1 | P97785 | 1 | 1 | -0.752 | -0.002 | 0.992 | 0.003 | 0.992 |
| Ttyh1 | Q9D3A9 | 2 | 2 | -0.752 | -0.002 | 0.988 | 0.005 | 0.992 |
| Serpina1d | Q00897 | 1 | 1 | 0.754 | 0.004 | 0.985 | 0.007 | 0.992 |
| Cntn4 | Q69Z26 | 5 | 5 | 0.755 | 0.005 | 0.980 | 0.009 | 0.992 |
| Nrcam | Q810U4 | 10 | 11 | 0.756 | 0.006 | 0.963 | 0.017 | 0.991 |
| Insr | P15208 | 1 | 1 | 0.756 | 0.007 | 0.976 | 0.011 | 0.992 |
| Grm3 | Q9QYS2 | 3 | 3 | 0.785 | 0.040 | 0.784 | 0.106 | 0.858 |
| Pecam1 | Q08481 | 1 | 1 | 0.792 | 0.047 | 0.762 | 0.118 | 0.856 |
| Ntm | Q99PJ0 | 2 | 2 | 0.793 | 0.049 | 0.840 | 0.075 | 0.891 |
| Ece1 | Q4PZA2 | 1 | 1 | 0.795 | 0.051 | 0.841 | 0.075 | 0.891 |
| Slc15a2 | Q9ES07 | 1 | 1 | 0.802 | 0.060 | 0.758 | 0.120 | 0.856 |
| Dkk3 | Q9QUN9 | 2 | 2 | 0.802 | 0.060 | 0.806 | 0.094 | 0.873 |
| Lingo1 | Q9D1T0 | 3 | 3 | 0.837 | 0.099 | 0.587 | 0.231 | 0.690 |

| | | | | | | | | |
|---|---|---|---|---|---|---|---|---|
| Calr | P14211 | 1 | 1 | 0.841 | 0.103 | 0.731 | 0.136 | 0.838 |
| Slc26a6 | Q8CIW6 | 1 | 1 | 0.854 | 0.117 | 0.836 | 0.078 | 0.891 |
| Grik2 | P39087 | 3 | 3 | 0.884 | 0.149 | 0.290 | 0.537 | 0.396 |
| L1cam | P11627 | 14 | 15 | 0.884 | 0.149 | 0.403 | 0.395 | 0.513 |
| Tmem63c | Q8CBX0 | 1 | 1 | 0.895 | 0.162 | 0.307 | 0.512 | 0.411 |
| Scn1b | P97952 | 3 | 4 | 0.902 | 0.169 | 0.515 | 0.288 | 0.625 |
| Grm1 | P97772 | 3 | 3 | 0.905 | 0.172 | 0.238 | 0.623 | 0.340 |
| Cfi | Q61129 | 2 | 2 | 0.914 | 0.182 | 0.416 | 0.381 | 0.526 |
| Lnpep | Q8C129 | 3 | 4 | 0.917 | 0.185 | 0.213 | 0.672 | 0.314 |
| Vapa | Q9WV55 | 1 | 1 | 0.923 | 0.191 | 0.329 | 0.483 | 0.432 |
| Gpm6b | P35803 | 1 | 1 | 0.926 | 0.195 | 0.420 | 0.377 | 0.528 |
| Pld3 | O35405 | 2 | 2 | 0.942 | 0.212 | 0.465 | 0.332 | 0.575 |
| Fgfr1 | P16092 | 1 | 1 | 0.950 | 0.220 | 0.496 | 0.304 | 0.606 |
| Tfrc | Q62351 | 1 | 2 | 0.961 | 0.231 | 0.302 | 0.520 | 0.410 |
| Asah1 | Q9WV54 | 2 | 2 | 1.002 | 0.273 | 0.222 | 0.653 | 0.322 |
| Grin2a | P35436 | 2 | 2 | 1.009 | 0.280 | 0.274 | 0.563 | 0.376 |
| Cd63 | P41731 | 2 | 2 | 1.011 | 0.283 | 0.100 | 0.999 | 0.177 |
| Lrrtm1 | Q8K377 | 2 | 3 | 1.032 | 0.304 | 0.164 | 0.784 | 0.259 |
| Gpc5 | Q8CAL5 | 2 | 2 | 1.038 | 0.310 | 0.158 | 0.801 | 0.252 |
| Pdgfra | P26618 | 1 | 2 | 1.076 | 0.347 | 0.274 | 0.563 | 0.376 |
| Trpc6 | Q61143 | 1 | 1 | 1.085 | 0.356 | 0.153 | 0.815 | 0.247 |
| P2rx7 | Q9Z1M0 | 1 | 1 | 1.093 | 0.365 | 0.156 | 0.807 | 0.250 |
| Prom1 | O54990 | 2 | 2 | 1.097 | 0.369 | 0.019 | 1.724 | 0.048 |
| Cntn1 | P12960 | 8 | 8 | 1.100 | 0.371 | 0.067 | 1.176 | 0.131 |
| Adgrl1 | Q80TR1 | 2 | 3 | 1.103 | 0.374 | 0.098 | 1.010 | 0.174 |
| Bpnt2 | Q80V26 | 1 | 1 | 1.104 | 0.375 | 0.205 | 0.689 | 0.312 |
| Anpep | P97449 | 2 | 2 | 1.112 | 0.383 | 0.095 | 1.021 | 0.171 |
| Nectin1 | Q9JKF6 | 5 | 5 | 1.116 | 0.387 | 0.144 | 0.842 | 0.236 |
| Slc6a2 | O55192 | 1 | 1 | 1.129 | 0.399 | 0.189 | 0.723 | 0.290 |
| Plpp1 | Q61469 | 1 | 1 | 1.130 | 0.400 | 0.069 | 1.158 | 0.133 |
| Sez6l2 | Q4V9Z5 | 3 | 3 | 1.152 | 0.421 | 0.052 | 1.283 | 0.110 |
| Itgam | P05555 | 2 | 2 | 1.156 | 0.425 | 0.077 | 1.113 | 0.145 |
| Cadm1 | Q8R5M8 | 5 | 6 | 1.187 | 0.454 | 0.048 | 1.315 | 0.104 |
| Abca7 | Q91V24 | 1 | 1 | 1.198 | 0.464 | 0.086 | 1.066 | 0.156 |
| Asic1 | Q6NXK8 | 1 | 1 | 1.206 | 0.471 | 0.027 | 1.567 | 0.065 |
| Cspg5 | Q71M36 | 1 | 1 | 1.227 | 0.490 | 0.146 | 0.835 | 0.238 |
| Itfg1 | Q99KW9 | 1 | 1 | 1.254 | 0.515 | 0.132 | 0.881 | 0.221 |
| Scn8a | Q9WTU3 | 3 | 4 | 1.260 | 0.520 | 0.014 | 1.855 | 0.040 |
| Olfm1 | O88998 | 3 | 3 | 1.295 | 0.552 | 0.015 | 1.835 | 0.040 |
| Plxna1 | P70206 | 3 | 4 | 1.324 | 0.577 | 0.007 | 2.175 | 0.024 |
| Ctsb | P10605 | 1 | 1 | 1.330 | 0.582 | 0.003 | 2.496 | 0.014 |
| Adra2a | Q01338 | 1 | 2 | 1.336 | 0.587 | 0.079 | 1.105 | 0.146 |
| Adam23 | Q9R1V7 | 2 | 2 | 1.348 | 0.598 | 0.037 | 1.430 | 0.081 |
| Serpina3k | P07759 | 2 | 2 | 1.359 | 0.607 | 0.036 | 1.441 | 0.080 |
| Lama4 | P97927 | 2 | 2 | 1.385 | 0.629 | 0.008 | 2.089 | 0.027 |

| | | | | | | | | |
|---|---|---|---|---|---|---|---|---|
| Ncam2 | O35136 | 3 | 3 | 1.424 | 0.662 | 0.004 | 2.381 | 0.017 |
| Slc36a1 | Q8K4D3 | 1 | 1 | 1.439 | 0.674 | 0.056 | 1.252 | 0.114 |
| Negr1 | Q80Z24 | 2 | 2 | 1.451 | 0.684 | 0.032 | 1.502 | 0.073 |
| Abca1 | P41233 | 3 | 4 | 1.462 | 0.693 | 0.000 | 3.602 | 0.003 |
| Calu | O35887 | 1 | 1 | 1.495 | 0.720 | 0.007 | 2.155 | 0.024 |
| Slc23a2 | Q9EPR4 | 1 | 2 | 1.534 | 0.751 | 0.035 | 1.456 | 0.079 |
| Pla2g7 | Q60963 | 1 | 2 | 1.542 | 0.757 | 0.005 | 2.327 | 0.018 |
| Lum | P51885 | 1 | 1 | 1.543 | 0.757 | 0.016 | 1.802 | 0.042 |
| Cp | Q61147 | 1 | 1 | 1.549 | 0.763 | 0.030 | 1.520 | 0.071 |
| Tenm1 | Q9WTS4 | 1 | 1 | 1.587 | 0.792 | 0.053 | 1.274 | 0.111 |
| Gria1 | P23818 | 2 | 3 | 1.609 | 0.809 | 0.000 | 3.870 | 0.002 |
| Sdk2 | Q6V4S5 | 1 | 1 | 1.727 | 0.896 | 0.005 | 2.267 | 0.020 |
| App | P12023 | 1 | 1 | 1.729 | 0.898 | 0.005 | 2.300 | 0.018 |
| Neo1 | P97798 | 4 | 4 | 1.746 | 0.910 | 0.001 | 2.973 | 0.006 |
| Uggt1 | Q6P5E4 | 1 | 1 | 1.844 | 0.979 | 0.008 | 2.080 | 0.027 |
| Ncan | P55066 | 3 | 3 | 1.863 | 0.992 | 0.007 | 2.147 | 0.025 |
| Trpv2 | Q9WTR1 | 1 | 1 | 1.890 | 1.010 | 0.001 | 3.160 | 0.005 |
| Gabra3 | P26049 | 1 | 1 | 1.954 | 1.053 | 0.000 | 4.195 | 0.001 |
| Atp6ap1 | Q9R1Q9 | 1 | 1 | 1.958 | 1.055 | 0.012 | 1.910 | 0.035 |
| Lsamp | Q8BLK3 | 2 | 2 | 2.034 | 1.104 | 0.001 | 2.890 | 0.007 |
| Brinp2 | Q6DFY8 | 1 | 1 | 2.105 | 1.149 | 0.003 | 2.582 | 0.013 |
| Cpd | O89001 | 2 | 2 | 2.110 | 1.152 | 0.000 | 4.791 | 0.001 |
| Drd1 | Q61616 | 2 | 2 | 2.149 | 1.176 | 0.000 | 4.027 | 0.002 |
| Bsg | P18572 | 4 | 4 | 2.162 | 1.184 | 0.000 | 4.849 | 0.001 |
| Abca5 | Q8K448 | 1 | 1 | 2.291 | 1.260 | 0.001 | 2.841 | 0.007 |
| Gpaa1 | Q9WTK3 | 1 | 1 | 2.297 | 1.264 | 0.002 | 2.709 | 0.010 |
| Lrrc8b | Q5DU41 | 1 | 1 | 2.415 | 1.330 | 0.000 | 3.721 | 0.002 |
| Nup210 | Q9QY81 | 1 | 1 | 2.438 | 1.343 | 0.000 | 4.363 | 0.001 |
| Itih5 | Q8BJD1 | 1 | 1 | 2.822 | 1.540 | 0.000 | 3.372 | 0.004 |
| Ces1c | P23953 | 3 | 3 | 2.924 | 1.588 | 0.000 | 4.249 | 0.001 |
| Igf2r | Q07113 | 1 | 1 | 3.213 | 1.718 | 0.000 | 4.708 | 0.001 |
| Atg9a | Q68FE2 | 1 | 1 | 3.460 | 1.820 | 0.003 | 2.521 | 0.013 |
| Cpe | Q00493 | 2 | 2 | 4.007 | 2.024 | 0.000 | 5.446 | 0.000 |
